# A distributive germline restricts the spread of new mutations

**DOI:** 10.64898/2026.02.05.704067

**Authors:** Justin Scherer, Maa Ahema Gaisie, Sungjin Park, Brad Nelms

## Abstract

Genetic bottlenecks in the germline can amplify the impact of mutations, increasing the likelihood that new mutations are transmitted to multiple offspring. Here, we evaluate the timing and consequences of bottlenecks leading to maize pollen. By tracking transposon-induced mutations across tissues, we find that pollen derives from multiple early embryonic cell lineages, maintained by radial symmetries throughout development. In controlled crosses, offspring from bulk pollen or apical branches rarely share mutations, whereas lateral branches sample fewer lineages and produce offspring with a 6.2-fold increase in mutation sharing. Thus, the persistence of multiple, early-diverged cell lineages into the germline reduces mutation recurrence without altering the underlying mutation rate. Similar principles may apply in animals, where diverse mechanisms ensure that multiple cell lineages contribute to the germline.

## Introduction

Germline mutations provide the primary source of heritable genetic variation, shaping patterns of genetic diversity and evolution. When a mutation occurs in a germline progenitor, it can spread through mitotic divisions and contribute to multiple gametes, causing offspring to share the same new mutations (*1–7*). The transmission of mutations to multiple offspring (hereafter “mutation recurrence”) has several consequences: it elevates the risk of disease recurrence within families (*1–3*), contributes to age-related infertility (*8–10*), and reduces the number of distinct variants entering a population (**Fig. 1A**, right), thereby limiting the mutation supply available for adaptation (*11,12*).

**Figure 1.**
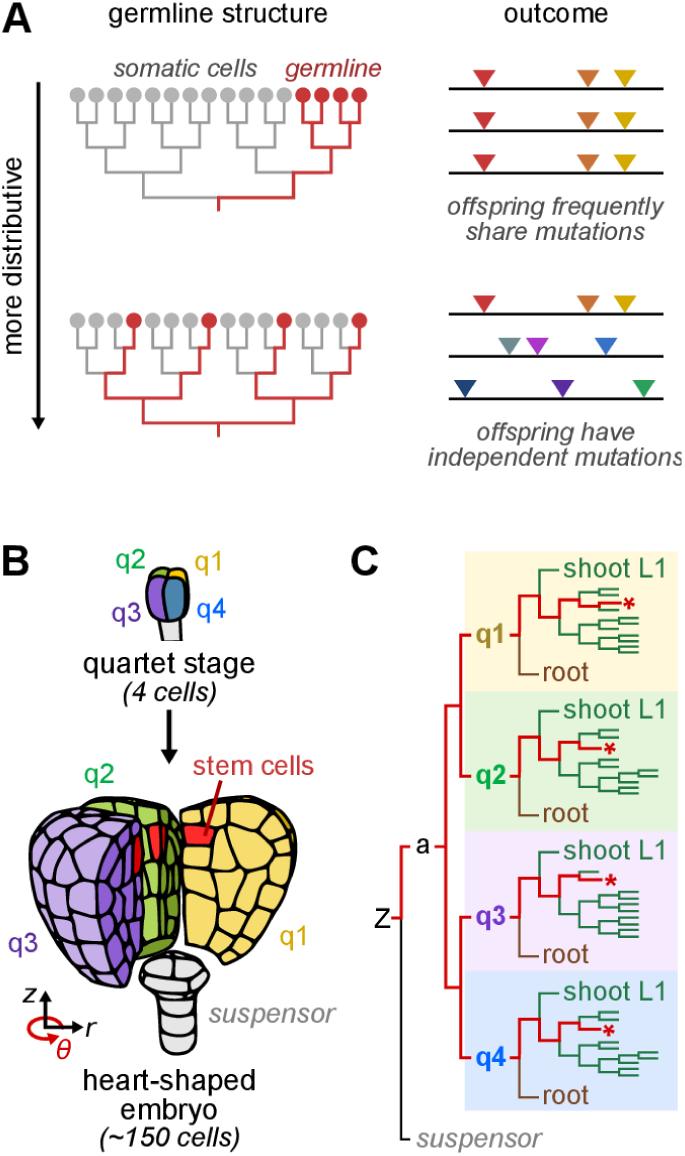
The distributive germline model. (**A**) Conceptual extremes of lineage architecture leading to the germline. If the germline were drawn from a single recent common ancestor (top), it would fix prior mutations and increase mutation recurrence in the offspring. Conversely, if the germline were drawn from multiple distant lineages (bottom), offspring would inherit independent mutations. (**B**) In *Arabidopsis*, the embryo has four symmetric quadrants, each descended from a distinct progenitor. Meristematic stem cell precursors, which ultimately contribute to the germline, arise at the immediate intersection of the quadrants. Embryo was segmented from microscope image in ref 23. (**C**) Lineage reconstruction for the embryo in panel B. Red asterisks, meristematic stem cell precursors.

Although many mechanisms regulate the germline mutation rate and spectrum (*13–16*), far less is known about the processes shaping mutation recurrence. In principle, mutation recurrence could be reduced in two ways. One would be to suppress mutations early in development, as late-arising mutations are less likely to contribute to multiple gametes. However, empirical support for this idea is limited: in humans, for example, the mutation rate is 3-fold higher during embryonic divisions compared to later in the germline (*3*).

An alternative strategy depends on how germ cells are related by descent during development. Because mutations are propagated through cell division, cell lineage relationships determine how widely mutations spread across an organism (*17–20*). If germ cells descend from a single recent progenitor (**Fig. 1A**), this creates a genetic bottleneck: mutations arising before the bottleneck would be transmitted to many gametes. At the other extreme, if germ cells descend from multiple, deeply diverged lineages (a “distributive germline”; **Fig. 1A**), then mutations would be shared only if they arose early in embryogenesis. Thus, by minimizing genetic bottlenecks in the germline, mutation recurrence could be reduced to a very low level, even without changes in the overall mutation rate.

Here, we track mutations caused by the active maize Mu transposon (*21*) across densely sampled pollen and matched somatic tissues. Using these data, we resolve the number, timing, and spatial distribution of cell lineages contributing to pollen. We then follow mutations into the offspring, directly connecting pollen lineage structure to *de novo* mutation transmission. Our results show that mutation rate and mutation recurrence are decoupled, demonstrating how lineage architecture shapes the rate of mutation recurrence among offspring.

## The distributive germline in plants

We first evaluated plant lineage tracing data (*17,22–27*) to assess when genetic bottlenecks might occur, or conversely be avoided, in the cells that give rise to the next generation. These data show that multiple, deeply diverged cell lineages may persist throughout the life cycle, driven by radial symmetries surrounding key stem and progenitor cell populations.

For example, in the *Arabidopsis* embryo, radial symmetries are established at the 4-cell (quartet) stage: each quartet cell forms a symmetric quadrant of the embryo, contributing to nearly all cell and tissue types (**Fig. 1B**). As a result, cells at the quadrant boundaries have distant lineage relationships, even when they are direct neighbors. Meristematic stem cell precursors, which ultimately give rise to the germline, form at the intersection of all four quadrants (*28*) (**Fig. 1B**), with each descended from a distinct quartet progenitor (**Fig. 1C**). This places their last common ancestor before many of the earliest recognized developmental milestones, including root-shoot axis establishment and separation of epidermal (L1) and subepidermal (L2/L3) layers. In maize, lineage-tracing supports the emergence of similar radially-partitioned lineages early in the embryo (*27*). Radial symmetries are then preserved in the meristem (**Fig. S1**), potentially maintaining multiple deep cell lineages throughout post-embryonic growth (*17*).

However, it remains unclear how robustly this lineage structure is preserved in practice. Meristematic stem cells form a small, positionally defined cell population that can undergo stochastic clonal replacement (*29,30*), raising longstanding questions about their permanence (*29*). Even a single clonal replacement event would fix early-arising mutations (*31,32*), erasing lineage diversity and negating the buffering effects of earlier radial symmetries. Determining whether multiple lineages persist into the germline therefore requires direct measurement of mutation spread through development and into the next generation.

## Spread of transposable element (TE) insertions through the maize tassel

In maize, pollen is produced by the tassel, a branched inflorescence composed of a central spike and 12-18 side branches (**Fig. 2A,B**). Each branch produces hundreds of thousands of pollen grains, which can be readily collected for sequencing or controlled crosses. By comparing the co-occurrence of mutations in pollen from different branches, we reasoned it would be possible to infer the spatial distribution and lineage relationships of mutant sectors within the tassel primordia. Importantly, side branches span a range of radial positions along the tassel circumference (**Fig. 2B**), where radial symmetries predict the strongest differences in lineage contribution and mutation spread.

**Figure 2.**
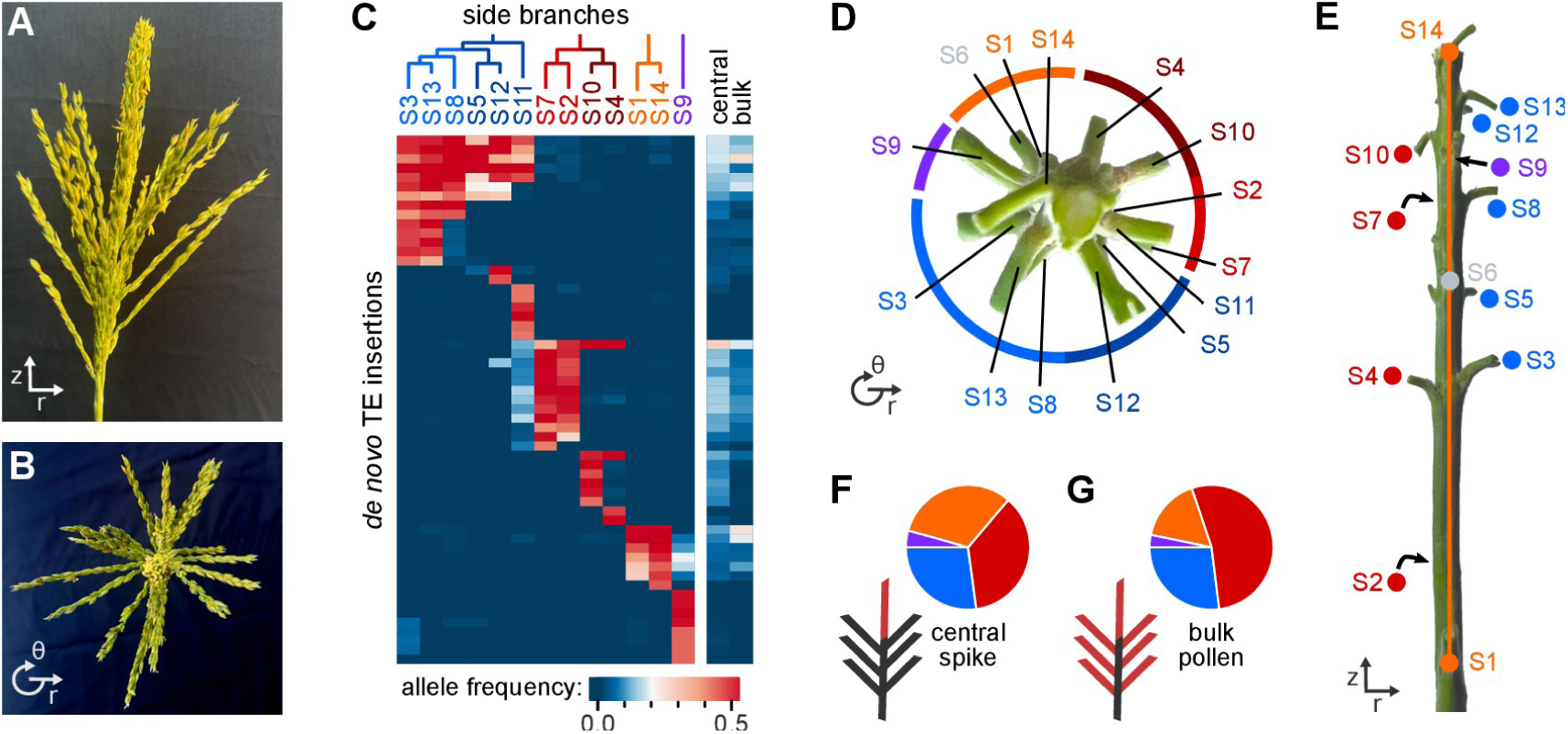
Multiple cell lineages contribute to pollen across the tassel circumference. (**A**) Side and (**B**) top view the maize tassel, showing the arrangement of branches. Each tassel contains 12-18 side branches, emerging from different circumferential positions (*θ*). (**C**) Allele frequencies of *de novo* TE insertions across pollen samples from a single plant. Allele frequencies range from 0 to 0.5, depending on the proportion of pollen grains with a given insertion. Top, maximum-parsimony lineage reconstructions for the cells giving rise to each side branch. (**D**) Side and (**E**) top view of tassel with branches cut back. Inferred lineages show strong correspondence with circumferential position (*θ*) rather than absolute distance. No data was obtained for branch S6, shown in gray. (**F,G**) Estimated fraction of central spike or bulk pollen derived from each color-coded lineage. Both central spike and bulk pollen contain a mix of lineages.

We collected pollen separately from multiple tassel branches in each of 11 Mu-active plants (**Fig. S2**) and measured the abundance of TE insertions by sequencing (*21*). To distinguish inherited from *de novo* TE insertions, we obtained matched endosperm tissue. The endosperm is derived from a sister sperm cell during double fertilization (**Fig. S3A**), and so TE insertions present in both pollen and endosperm must have occurred prior to the zygote. As expected, TE insertions present in the endosperm were found throughout the tassel (**Fig. S3B**), consistent with their identification as ubiquitous, inherited TEs. In contrast, *de novo* TE insertions (absent from endosperm) were found in only subsets of the tassel, present in pollen from one or a few branches (**Fig. 2C**).

The most abundant *de novo* insertions (allele frequency > 0.45) were often found in multiple branches of the same tassel (**Fig. 2C**), whereas no such sharing was observed between pollen from different plants (*p* < 10^-5^, permutation test; **Fig. S3B**). This sharing is unlikely to reflect cross-contamination: there were consistently multiple high-abundance insertions uniquely marking each branch, whereas experimentally mixed samples showed correlated abundances across all insertions (**Fig. S4**). Instead, shared *de novo* insertions are best explained by a single transposition event that occurred early in development and subsequently spread via clonal expansion.

## Multiple cell lineages contribute to pollen across the tassel circumference

If early transposition events are spread clonally, their spatial distribution should reflect the underlying lineage structure of the tassel. Indeed, side branches from the same plant clustered according to shared *de novo* TE insertions (**Fig. 2C, S5-S13**): branches within a cluster shared multiple insertions, while branches in different clusters often shared none. Using the presence and absence of *de novo* insertions, we reconstructed maximum-parsimony phylogenies for the cell lineages contributing to pollen in each branch (**Fig. 2C**, top; **Supplementary Note 1**).

Mapping the inferred cell lineages onto the physical positions of tassel branches revealed a striking spatial organization (**Fig. 2D, E**). Lineages were closely associated with position along the tassel circumference when viewed from above (**Fig. 2D**), consistent with radial symmetries established during embryo and meristem development (*17,27–29*) (**Fig. 1**). Branches separated by large vertical distances but aligned along the circumference were frequently derived from the same lineage (e.g. branch ‘S1’ and ‘S14’; **Fig. 2E**). Conversely, adjacent branches often belonged to different lineages when they were separated along the tassel circumference (e.g. ‘S8’ and ‘S7’; **Fig. 2D**). Finer-scale alignment along the circumference predicted sublineage structure, with closely aligned branches sharing the greatest overlap of *de novo* insertions (**Fig. 2C,D**).

This lineage structure directly shaped mutation abundance in pollen. A single side branch samples less than 10% of the tassel circumference and therefore was often derived from one or few early clonal lineages. Accordingly, an average of 3.8 *de novo* TE insertions were mitotically fixed per side branch (allele frequency = 0.5; **Fig. 2C**). In contrast, the central spike forms near the meristem apex and integrates contributions from multiple lineages, resulting in <1 fixed *de novo* insertions per branch. Consistent with this interpretation, pollen from the central spike closely resembled bulk pollen collected from a mixture of branches (**Fig. 2C**, right). Both sources integrate lineage diversity across the tassel circumference (**Fig. 2F,G**), preventing any single *de novo* insertion from reaching high abundance.

## Embryonic origin of pollen lineages

To establish when pollen lineages emerge during embryogenesis, we sequenced matched leaf tissue from seven plants. In maize, leaves emerge sequentially from the meristem, and clonal sectoring studies have established the timing at which leaf progenitors diverge from pollen (*25–27*). By seed maturity, all leaves have diverged from the tassel central spike (*25,26*): 4-6 leaf primordia are already formed in the embryo, while later leaves arise from meristematic cells basal to the tassel progenitors. Consequently, TE insertions shared between pollen and leaves can be traced to embryonic development.

Each plant contained multiple *de novo* TE insertions shared by pollen and leaves (11.6 ± 2.3 per plant), indicating that the Mu transposon is active during seed development and can mark cell lineages within the embryo. We used the earliest leaf in which a TE insertion was detected as a proxy for developmental timing (**Supplementary Note 1**). **Fig. 3A** shows a phylogenetic reconstruction for the embryonic cell lineages that gave rise to pollen for the plant in **Fig. 2**. Because lineages can only be detected after being marked by an insertion, some ambiguity is unavoidable, resulting in multiple equally parsimonious trees (e.g. the relationship between the 4 earliest cell lineages in **Fig. 3A** is uncertain). Nonetheless, the number of inferred lineages at each stage provides both an upper and lower bound for how many embryonic cells contributed to pollen over time (**Supplementary Note 2**).

**Figure 3.**
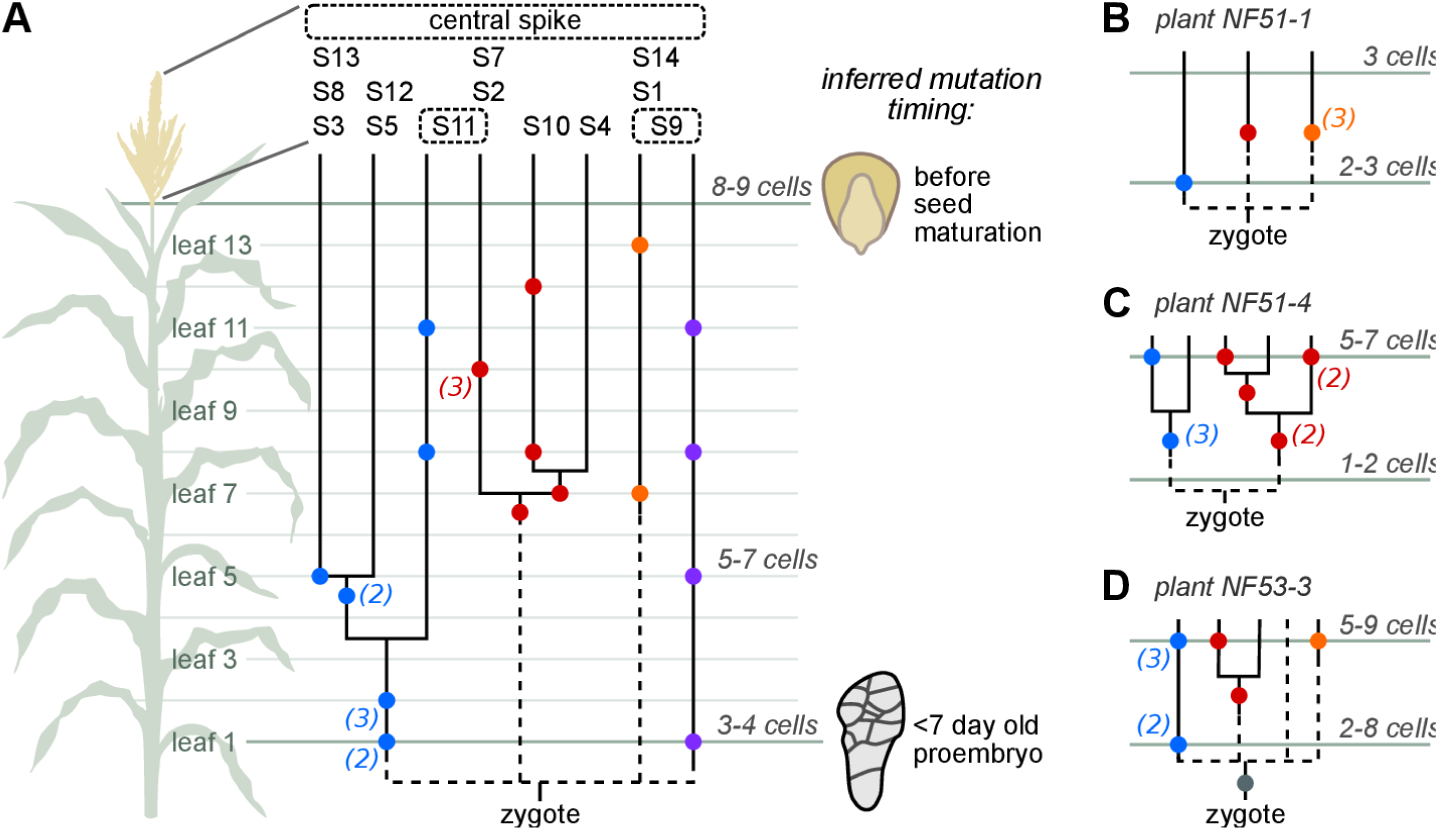
Pollen lineages originate early in the embryo. (**A**) Phylogenetic reconstruction of lineages contributing to pollen for the plant in Fig. 2. Colored dots, TE insertion(s) detected in both pollen and leaves; when multiple insertions were detected in the same set of samples, the number of insertions is shown in parentheses. Vertical positions of dots indicates the first leaf in which a given insertion was detected. Inferred mutation timing based on sectoring studies (*25–27*). (**B-D**) Lineage reconstructions for three additional plants. Bottom horizontal line, insertions detected in the first seedling leaf; top horizontal line, insertions detected in the topmost leaf. Complete lineage trees and supporting data are in **Fig. S7-S13**.

Using this framework, we find that the number of pollen progenitors increases from approximately 2–3 at the time of first leaf divergence to 5–6 in the mature embryo (**Fig. 3, S14**). The relative contributions of early progenitors to the final pollen population was variable. In some cases, lineage abundance was largely preserved over time (e.g., one of two early progenitors contributed to 50% of pollen), whereas in others, early lineages were substantially diluted or expanded. For example, the purple lineage in **Fig. 3A** initially represented one of three or four pollen progenitors, but ultimately contributed to only ∼5% of total pollen (**Fig. 2F,G**). This variability supports the view that pollen progenitor contributions are shaped by stochastic lineage dynamics and positional effects within the meristem (*27*), and do not reflect rigidly determined cell fates.

Despite this stochasticity, our results place the origin of pollen lineages among the earliest embryonic divisions. In X-ray sectoring experiments (*27*), mutant sectors that extend from the first leaf into the tassel were only observed when X-rays were applied during the first 4 days of development (the 4-20 cell proembryo), never later. Five plants were marked by *de novo* insertions in both pollen and the first leaf, and, in 3 of these (60%), there must be at least two cell lineages to explain the phylogeny (**Fig. S14A**). Thus, multiple germinal lineages are established early in embryogenesis and persist throughout development.

## Pollen lineages preserve genetic diversity among the offspring

If pollen derives from multiple, spatially segregated lineages, this organization should directly influence how new mutations are transmitted to offspring. To test this, we performed controlled crosses (**Fig. 4A**) using pollen collected from either the full tassel (“bulk” pollen), an individual side branch, or the central spike. A Mu-inactive line served as the female parent, isolating any new Mu activity to the male. For each cross, we collected endosperm from the Mu-active parent to identify ancestral TE insertions, and sequenced both endosperm and leaf tissue from 5-10 offspring.

**Figure 4.**
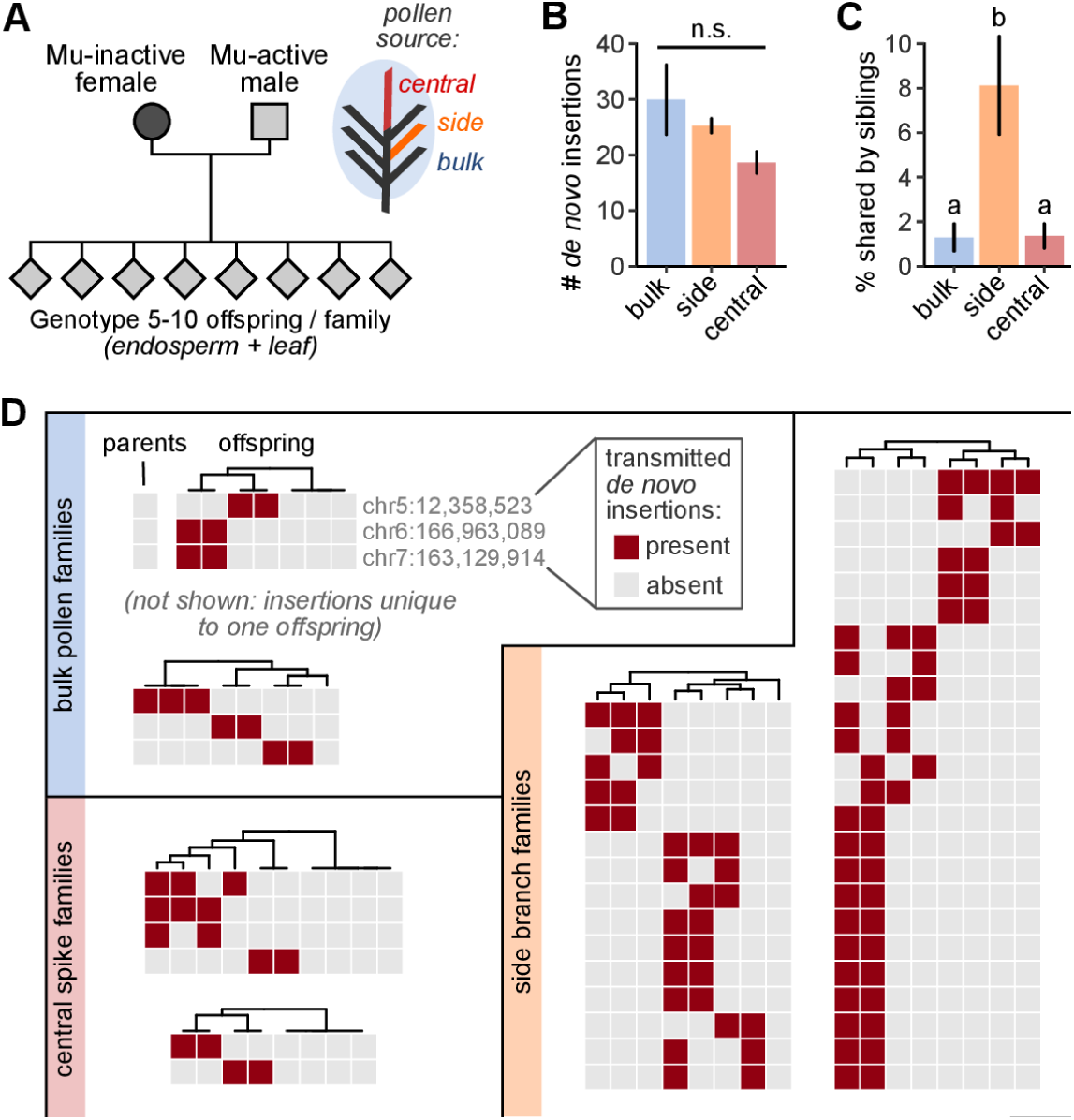
Mutation recurrence among the offspring is sensitive to pollen source. (**A**) Experimental design. Controlled crosses were performed with pollen collected in bulk, from the central spike, or single side branches. (**B**) The mutation rate does not differ significantly by pollen source. (**C**) Mutation recurrence is significantly higher in crosses using side branch pollen (Tukey’s Honest Significant Differences test). (**D**) Two families of each cross type, showing clustered mutation sharing from side-branch pollen versus largely independent mutation sets from bulk or central spike pollen.

On average, 26.0 *de novo* TE insertions were transmitted per offspring, with no significant differences between crosses using bulk, side branch, or central spike pollen (**Fig. 4B**). Thus, pollen source had little effect on the the overall mutation rate, consistent with prior results (*7*). In contrast, mutation recurrence differed significantly (**Fig. 4C**). Offspring from side-branch pollen shared *de novo* insertions 6.2-times more often than offspring from bulk or central spike pollen. Moreover, side branch offspring frequently shared large clusters of insertions (**Fig. 4D**), while such clusters were rare among offspring derived from bulk or central spike pollen.

These differences mirror the underlying lineage structure of the pollen source: side branches typically sample a single embryonic lineage, whereas bulk and central-spike pollen integrate contributions from multiple. Thus, the spatial origin of pollen – rather than the mutation rate itself – dominates the probability of recurrent mutation transmission among offspring. By sampling from multiple, deeply diverged cell lineages across the circumference, the maize tassel preserves genetic diversity among offspring and limits mutation recurrence within families.

## The distributive germline in animals

We next examined existing lineage and developmental data to ask whether animals might also maintain multiple, deeply diverged germline lineages, focusing on humans and four well-studied model organisms: mouse, *Drosophila*, *C. elegans*, and zebrafish. Unlike plants, all five animal species set aside a dedicated germline early, with progenitor germ cells (PGCs) among the first cell types to be specified in the embryo. However, early germline specification does not necessarily imply that the germline is derived from a single clonal lineage. Indeed, with the exception of *C. elegans*, germline progenitors in these species are polyphyletic (**Table S1**), with subsets of germ cells more closely related to somatic lineages than to each other (*1–3,33–37*).

A range of developmental mechanisms ensure that multiple cell lineages contribute to animal germlines. In zebrafish, the germline is specified by maternally deposited germ granules. These granules aggregate along the first four division planes and then segregate asymmetrically to one daughter cell throughout the next 8 divisions (*33*) (**Fig. 5A**). This process ensures that each of the first four embryonic cells contributes deterministically to one quarter of the germline.

**Figure 5.**
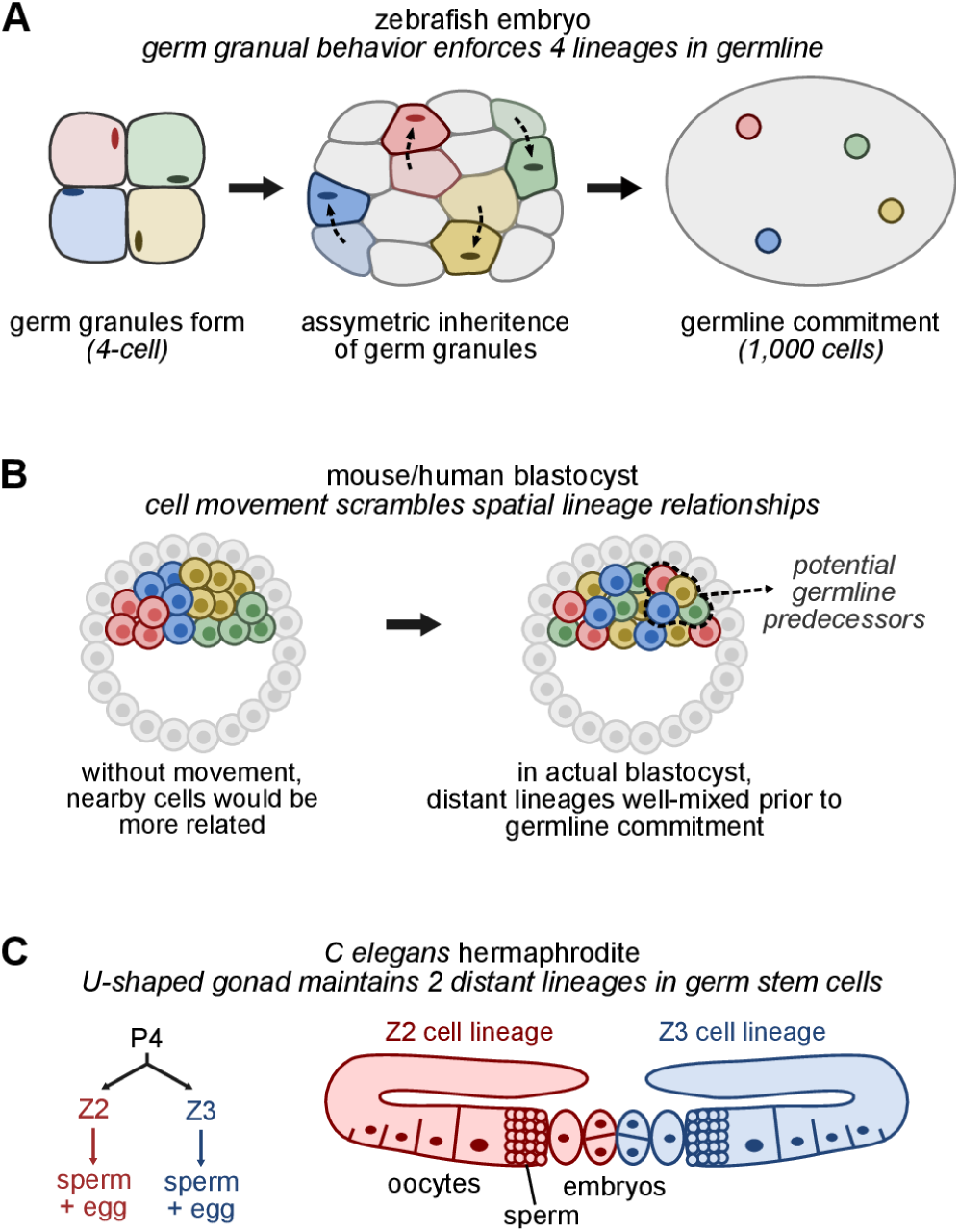
Diverse mechanisms maintain multiple, early-diverged cell lineages in animal germlines. (**A**) Zebrafish germ granules form at the 4-cell stage and then segregate asymmetrically to one daughter cell during subsequent divisions. Progenitor germ cells are later specified based on the presence of germ granules, with each descended from a distinct 4-cell initial. (**B**) In mammals, early intermixing of cells disrupts spatial lineage continuity and randomizes the contribution of early embryonic lineages to the germline. (**C**) In *C. elegans*, the germline progenitor cell (P4) divides into two germ cell initials (Z2, Z3). Each initial then forms a separate gonad arm, restricting the spread of mutations arising in either lineage to one half of the gonad.

In mammals, pre-gastrulation embryos undergo rapid cell movements (*34,35*), disrupting lineage relationships between neighboring cells (**Fig. 5B**). As a result, progenitor germ cells typically arise from multiple lineages (*37*). In humans, *de novo* mutations that are transmitted to offspring can be detectable in subsets of parental blood (*1,3*), indicating that multiple early cell lineages contribute to both soma and germline.

Even in *C. elegans*, which has a single post-zygotic germline progenitor (P4), the adult hermaphrodite gonad comprises two arms derived from different germline initials (Z2 and Z3; **Fig. 5C**). Germline stem cells sometimes undergo clonal replacement within a gonad arm (*32*), and so this U-shaped structure limits the transmission of mutations arising in either gonadal lineage to at most half of the germ cells. Thus, across species, germline specification mechanisms that appear complex from a cell-fate perspective can instead be understood as strategies that limit early fixation of new mutations by preserving lineage diversity.

## Perspective

Our results show that cell lineage architecture leading to the germline plays a central role in shaping how often mutations are shared among offspring. Both plants and animals show features of a distributive germline, in which germinal precursors are derived from multiple, deeply diverged cell lineages. This architecture limits genetic bottlenecks and reduces mutation recurrence among the offspring, even when mutations arise early.

Other mechanisms have been proposed to limit mutation accumulation in plants, particularly within the meristem (*17,29,30,38*). For example, selection among meristematic stem cells could purge deleterious new mutations (*30,38*); while limited selection could occur within a distributive germline, the two strategies are in tension: sustained intra-organism selection necessarily reduces lineage diversity and would increase mutation recurrence among offspring. Beyond selection models, Burian and colleagues (*17*) recognized that cell division patterning within the meristem may preserve genetic diversity in long-lived plants, providing an important conceptual predecessor to the distributive germline framework we develop and test here.

Overall, our findings highlight mutation recurrence as a developmental and evolutionary parameter distinct from mutation rate itself. The repeated emergence of distributive germline architectures in plants and animals suggests that buffering against early mutational fixation may be broadly advantageous. Determining when such buffering is favored – versus when bottlenecks (*39*) or stem cell competition (*30,38*) are beneficial – will require theory that integrates development, lineage structure, and selection across life histories.

## Acknowledgments

We thank Jonathan Gent, Bob Schmitz, Zack Lewis, Max Staller, and Kelly Dawe for invaluable discussions and critical reading of the manuscript.

## Author Contributions

Design and conception, B.N.; Data collection, J.S. and S.P.; analysis, J.S., M.A.G., B.N.; Writing, B.N. with input from other authors.

## Data and materials availability

Sequencing data will be deposited to the Gene Expression Omnibus and made public after peer review.

## Materials and Methods

### Segmentation and lineage reconstruction of an early *Arabidopsis* embryo

For the *Arabidopsis* embryo in **Fig. 1**, a Z-stack microscope image of a heart shaped embryo was downloaded from ref. 40. The embryo was segmented using the MorphoLibJ package (*41*), reproducing the procedure of ref 23. Cell lineage relationships were then reconstructed based on the position of cell division planes using the TreeJ package (*23*). Lineage trees were constrained to converge on the known *Arabidopsis* embryo lineage relationships at the 64-cell stage, as determined by live-cell microscopy (*22*).

### Plant growth and tissue collections

The Mu-active line was descended from maize Co-op stock 919J and maintained by continual outcrossing onto Mu-inactive Co-op stock 910I, as described (*21*). Before planting, all seeds were incubated in water for 10 min and then chipped with a razor blade to collect endosperm tissue; endosperm samples were stored at -80 °C until further processing. Seeds were then planted in the Botany Greenhouses in Athens, Georgia and grown in sunlight supplemented with LED fluorescent lights (Medic Grow 550W Slim Power 2). For leaf collections, a ∼½-inch (for seedling leaves) or ∼2-inch (for adult leaves) piece of the leaf was cut and stored frozen. Pollen was harvested at maturity, using shoot bags (Seedburo Equipment Company cat. no. S27) to collect pollen from different branches (**Fig. S2B**) or tassel bags to collect bulk pollen from the entire tassel.

### Mu TE library preparation and sequencing

DNA extractions and MuSeq2 library preparation were performed as described (*21*). Briefly, frozen samples were first ground to a fine powder, using a mortar and pestle for endosperm and leaf and a Qiagen tissue lyser II (Qiagen cat. no. 85300) for pollen. DNA was extracted using the Qiagen Dneasy Plant Mini kit (Qiagen cat. no. 69104) for leaf, and Cetrimonium bromide (CTAB) for endosperm and pollen. DNA was then sheared with a Covaris E220 Evolution Sonicator (Covaris cat. no. 500429) to a final average size of ∼350 bp (for the maize tassel maps in **Fig. S5-S11**) or 1000 bp (for all other samples). For pollen samples, a second round of DNA cleanup was performed after shearing using Monarch DNA spin columns (New England Biolabs cat. no. T2038L).

DNA was end-repaired and MuSeq2 adapters (*21*) were ligated to the ends using the NEB Ultra II DNA Library Prep Kit for Illumina (New England Biolabs cat. no. E7645). Pollen samples were prepared using ½ the normal volumes from the NEB kit; endosperm and leaf were prepared using volumes further reduced to 1/8 of a standard reaction. Adapter-ligated DNA was purified and concentrated using AmpureXP beads (Beckman Coulter), and Mu TE insertions were amplified through a series of three nested PCR reactions as described (*21*). Libraries were sequenced by Admera Health on a NovaSeq X Plus, using paired-end 150 bp reads and 10% PhiX added.

### TE insertion mapping and analysis

TE insertions were mapped to the maize W22 genome (*42*) and quantified as described (*21*). MuSeq2 libraries use unique molecular identifiers (*43*) to identify PCR duplicates and infer the number of DNA molecules that were sequenced from the original (unamplified) sample. After mapping, ‘historical insertions’ – a set of 29 insertions present in the Mu-inactive maintainer line (21) – and a previously defined set of 19 ‘blacklisted’ sites (21) were removed. The blacklist sites represent locations we consistently find at a low level across all libraries; many of these show weak homology to the Mu terminal inverted repeats in the reference genome and may represent diverged, ancestral TE insertions.

For all plants, the Mu-inactive line was used as the female parent. This line contains a static set of 29 historical insertions, and so other insertions were either inherited from the male parent or arose *de novo* in the individual. To distinguish paternally inherited from *de novo* insertions, we used matched endosperm tissue: insertions above 100 counts per million (CPM) in both endosperm and another tissue (e.g. leaf or pollen) were classified as paternally inherited. There was a clear separation of inherited from *de novo* insertions using this approach, with very little sensitivity to the chosen threshold (see ref. 21 and **Fig. S3B**).

Paternally inherited insertions were then used to normalize relative molecule counts (e.g. CPM) to absolute allele frequencies in the underlying sample. The Mu transposon has a very low excision rate in pollen (∼10^-4^ per generation; *44*), and so paternal insertions (absent from the mother) will be at an allele frequency of ∼0.5 in all samples (heterozygous). We used the known abundance of paternal insertions to estimate unknown allele frequencies for *de novo* insertions: the relative molecule abundances were rescaled so that the average paternal insertion has an allele frequency of 0.5. The accuracy of this approach is discussed and validated in greater detail in ref. 21.

For the tassel sectoring data (e.g. **Fig. 2** and **3**), samples were required to pass two quality control criteria: (i) at least 5000 molecules were sequenced spanning the TE-genome junction and (ii) the normalization error for converting molecule counts to allele frequencies was <10%. Normalization error was estimated as the standard error of the mean for the average paternal insertion abundance. This second criterion removed a subset of samples where paternal TE insertions were inconsistent in abundance; we had not seen this behavior before with MuSeq2 samples (*21*), but for a subset of side branches with very low pollen release the data were noisier and the normalization error criterion removed such samples. One sample from plant NF53-2 – side branch 5 (‘S5’) – failed these normalization criteria but was featured in the main text anyway; this sample was shown as the data was sufficiently clean to define this branch within the ‘blue’ cluster, and we favored having more complete information on the tassel branches when plotting the sector distributions.

### Lineage reconstruction

To infer pollen lineage phylogenies from the Mu TE insertion data, we focused on *de novo* TE insertions that surpassed an allele frequency of 0.45 in at least one pollen sample. The rationale for 0.45 is that it is nearly fixed within the diploid progenitor cells (0.5 would be fixed) but allows for slight measurement error. The number of TE insertions to analyze (e.g. how many insertions were in a heatmap such as **Fig. 2C**) was sensitive to this threshold choice, however the lineage reconstruction was robust to lowering this threshold – insertions below the cutoff consistently supported the same branch relationships as those above it. The process and rationale used to infer maximum parsimony phylogenies from the TE insertion data are described in **Supplementary Note 1**.

### Analysis of TE data from outcross experiments (**Fig. 4**)

For each outcross family, TE insertions were sequenced in both chipped endosperm and matched tissue derived from the embryo (typically the first seedling leaf). Inherited insertions were then called as having ≥500 CPM in either leaf or endosperm and ≥200 CPM in the other. To distinguish de novo insertions that arose in the male parent but then were transmitted to the offspring, we further excluded any insertions detected at ≥200 CPM in the parental endosperm. After identifying these transmitted *de novo* insertions, we calculated the average number of transmitted insertions in each family as well as the average % shared by siblings. **Fig. 4BC** shows a boxplot of the family averages, with N = 6 families derived from bulk pollen, 5 from side branch pollen, and 3 from central spike pollen. As each family-wide measurement required data from multiple tissues and individuals, a total of 228 tissue samples from 121 plants were analyzed for this experiment.

### Source images used for figure cartoons

To increase realism, several main text cartoons depicting different developmental stages were drawn by hand or in Inkscape using published microscope images as a reference. The *Arabidopsis* embryo in **Fig. 1** was created by segmenting a supplemental Z-stack image from ref. 23 (as described above). The maize embryo in Fig. 3A was drawn from micrographs in ref. 27 and depicts the latest embryonic stage in which sectors spanning from the first leaf to the tassel were observed in that study. The 16-cell zebrafish embryo in Fig. 5A was drawn from an image in ref. 45.

### Use of generative AI

ChatGPT-5 was used to assist with text editing. We found it gave valuable suggestions to clarify the writing. However, the AI could not reliably identify changes it had made and sometimes produced incorrect statements. To address this limitation, all writing and editing was maintained in an offline document. AI-suggested changes were then assessed by the authors and manually implemented if they were deemed useful. This process ensured that human input was present at every step of the writing.

## SUPPLEMENTARY MATERIALS

**Fig. S1.**
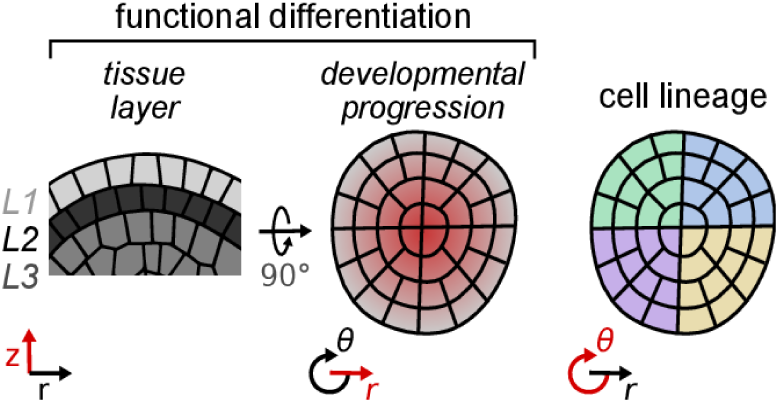
Radial symmetries may preserve multiple cell lineages in the plant meristem. Side (left) and top (middle, right) views of the shoot apical meristem (SAM), labeled by (left) tissue layer, (middle) developmental progression (e.g. stem cell vs more differentiated), and (right) cell lineage. These cartoons are based on *Arabidopsis*; similar symmetries exist in maize, but there are not separate L2 and L3 layers, instead the L2 has divisions occurring across 3 dimensions rather than forming a single (tunica) layer. The SAM is organized with a small cluster of stem cells in a region called the central zone (CZ). When these cells divide, their progeny are pushed outward to initiate organ primordia. If the CZ were reliably centered around the same cells, it would maintain multiple cell lineages throughout development (*17*). Essentially, plants organize their embryos (Fig. 1B) and meristems (this figure) so that function and lineage are specified by orthogonal coordinates: functional differentiation occurs radially (*r*) and axially (*z*), while lineages are maintained along the circumference (*θ*).

**Fig. S2.**
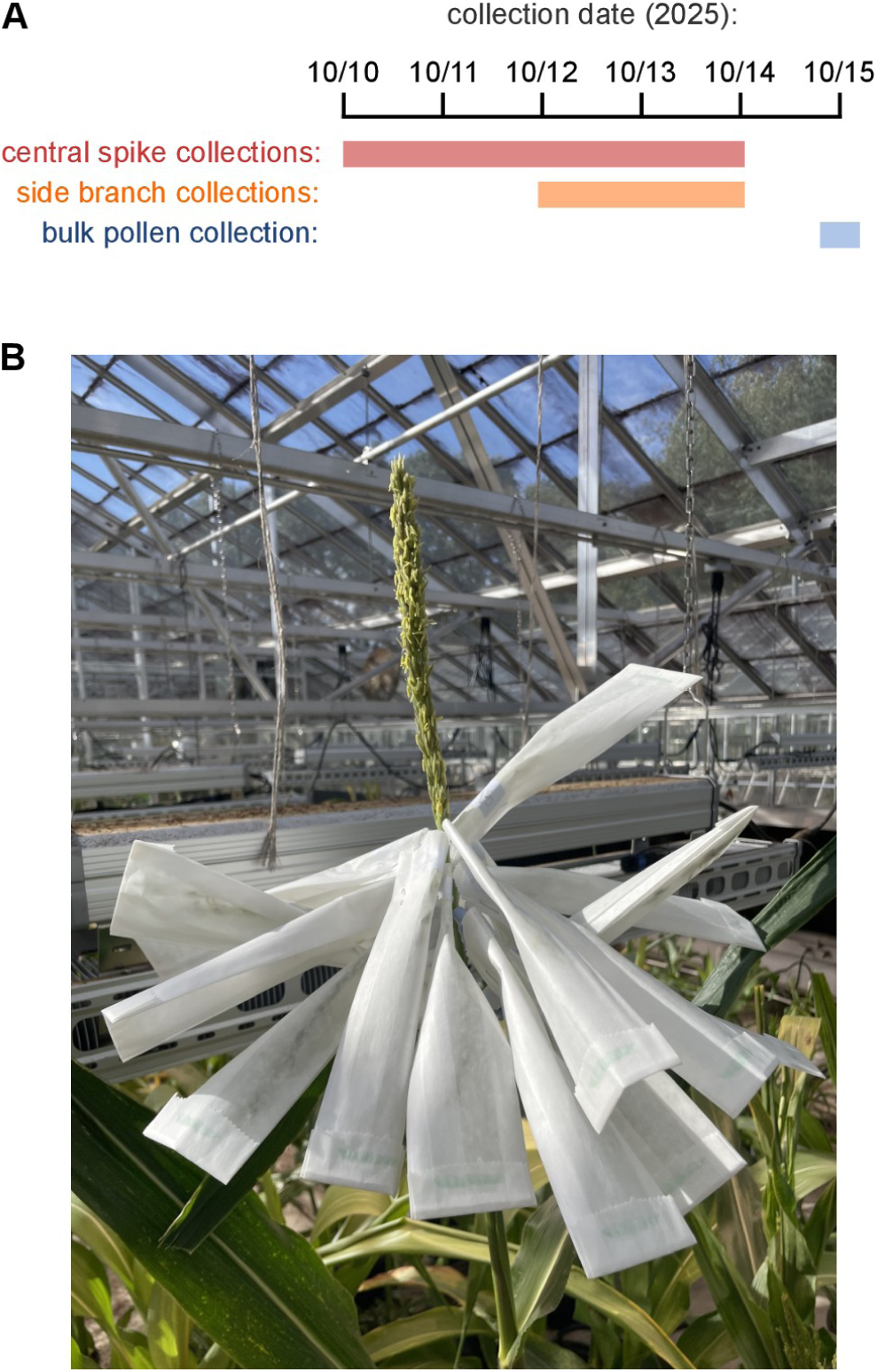
Experimental design for tassel collections. **(A)** Time-course of pollen collections for a representative plant (plant NF53-2, the plant shown in **Fig. 2** and **3**). Multiple central spike collections were performed on consecutive days while pollen was shedding. Side branches generally start shedding later and were collected using individual bags. After all side branch collections, a single bulk pollen collection was performed for a subset of plants by placing a bag over the entire tassel the next day. **(B)** Picture of a tassel with bags placed on side branches for collections.

**Fig. S3.**
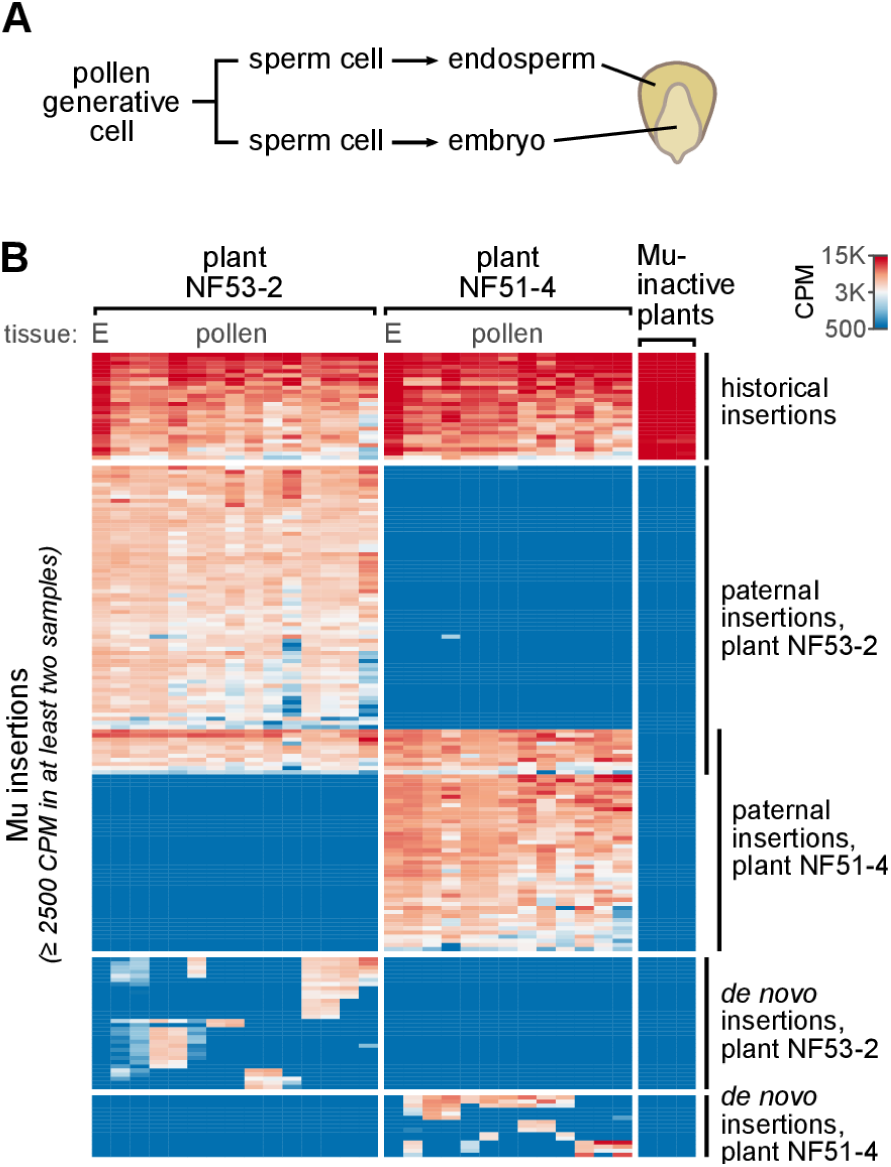
Distinguishing inherited from *de novo* TE insertions using matched endosperm. (A) A single mitotic division in pollen generates two sister sperm cells. These provide the paternal contributions to the embryo and endosperm during double fertilization. As a result, the paternal endosperm contribution diverges from the embryo precisely one cell division before the zygote. (B) Heatmap of abundant insertions in two representative Mu-active plants plus the Mu-inactive line used in this study. The tissue source is labeled on top; for the Mu-active plants, the first bar (E) shows the endosperm data and the remainder show pollen collections from different tassel branches. All insertions detected at an abundance > 2500 counts per million (CPM) are shown, using the same pipeline as was used to generate **Fig. 1** of ref 21. TE insertions are divided into categories based on the presence in different samples: historical insertions are present in the Mu-inactive line, paternally inherited insertions are present in endosperm as well as all tissue samples from a plant (we know these are paternally inherited because the Mu-inactive line was used as the female parent throughout this study), while *de novo* insertions are absent from matched endosperm (bottom).

**Fig. S4.**
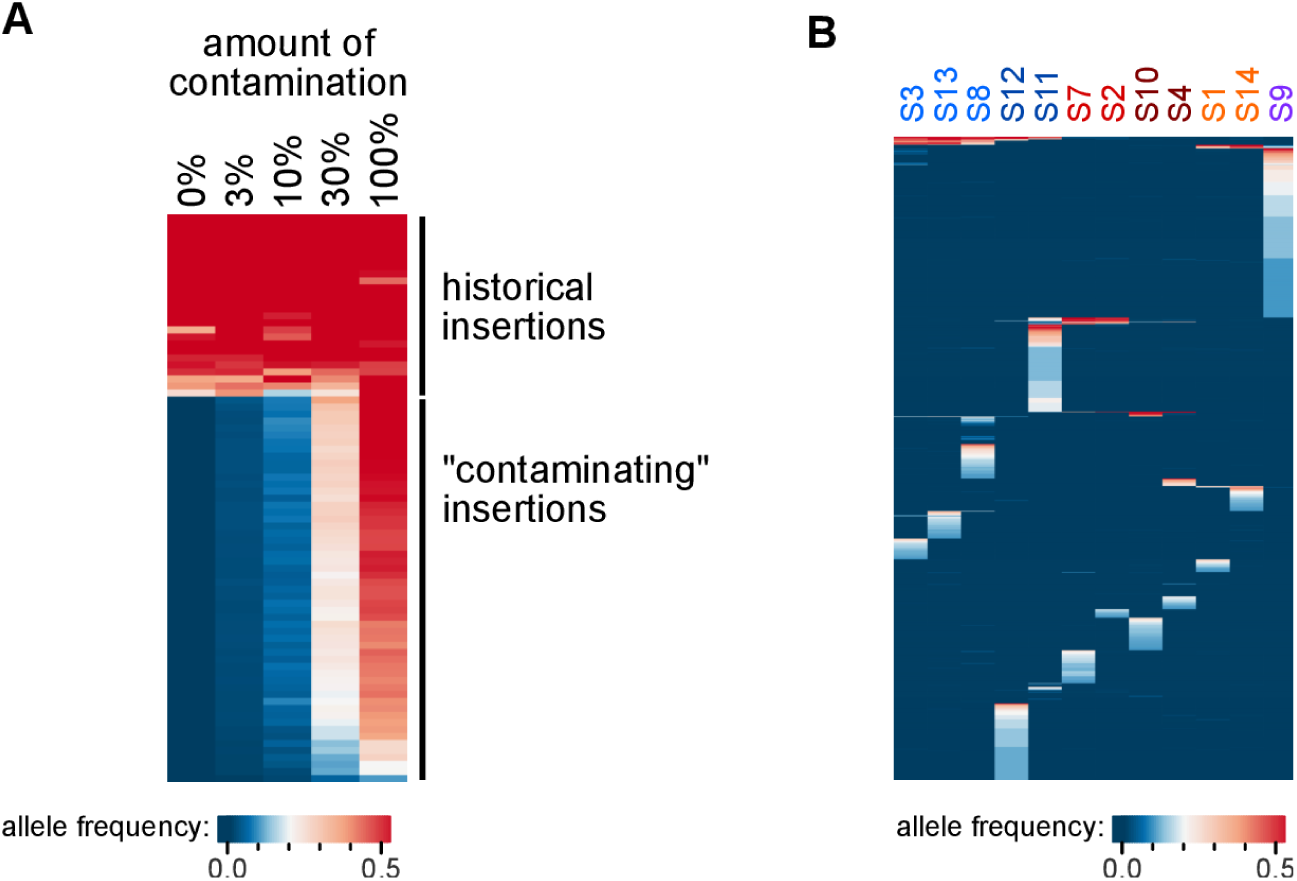
*De novo* TE insertions found in multiple tassel branches cannot be explained by cross-contamination. (A) This panel shows an example of intentionally mixed samples, mimicking how cross-contamination might appear in MuSeq2 data. DNA from two samples were experimentally mixed at defined ratios and then MuSeq2 libraries were prepared from each. One of the samples (the “non-contaminating” sample) is from a Mu-inactive plant, and has a defined set of 29 historical insertions. The other “contaminating” sample is from a Mu-active plant and was mixed to be 3%, 10%, 30%, or 100% of the total DNA. All insertions with an allele frequency > 0.1 in at least one sample are shown. When contamination is present, all insertions from the contaminating sample were detected and the measured allele frequency was proportional to the amount of contamination. Data re-analyzed from ref. 21. One feature of the tassel branch data that rules out cross-contamination is that, in the actual side branch data (e.g. Fig. 2C), the measured allele frequency for insertions shared by side branches is very high (>0.45). Such high abundances were not observed in artificially mixed samples, even when the contaminating sample was mixed as a high proportion of the total (30%). (B) A second feature that rules out cross-contamination is that every side branch had multiple insertions unique to it and not present among other samples. This heatmap shows measured de novo TE allele frequencies for side branches from the plant shown in **Fig. 2C**, showing any *de novo* insertions detected at an allele frequency > 0.1 in at least one sample (**Fig. 2C** was similar but used a much higher allele frequency cutoff of 0.45). Artificially mixed samples (e.g. panel **A**) do not show such unique insertions; rather, all insertions present in the contaminating sample were detected.

**Fig. S5.**
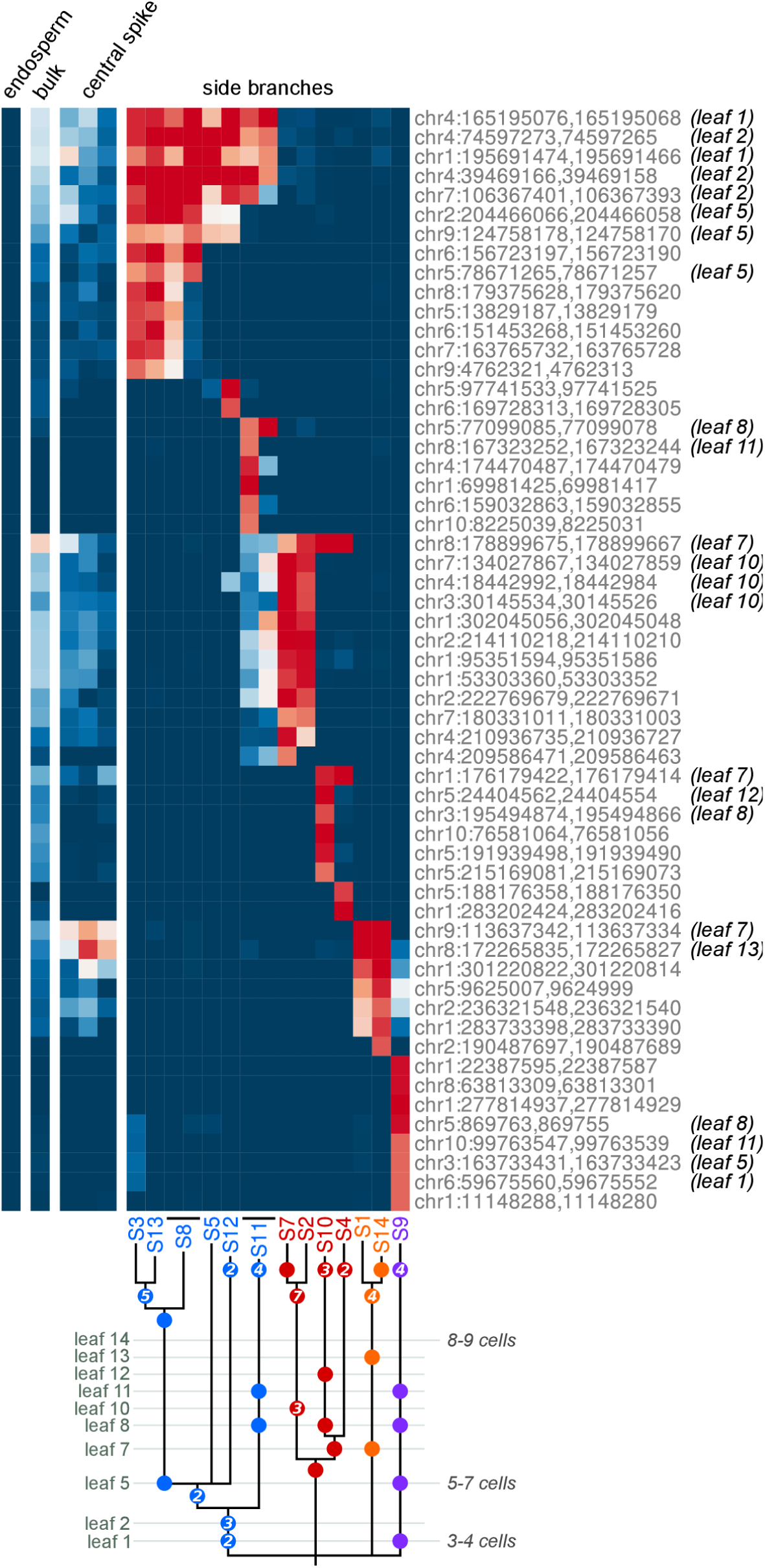
Heatmap of abundant *de novo* insertions and lineage reconstruction for plant NF53-2. This is the same plant shown in **Fig. 2** and **3** of the main text. Top, the heatmap from **Fig. 2C** is reproduced with additional information: (i) the chromosomal location of each insertion is labeled, (ii) pollen was collected multiple times from a subset of branches on subsequent days – these repeat collections were left out of the main text for simplicity but are included here, (iii) insertions detected in leaves were specifically labeled. For branches with repeat collections, samples from the same branch were placed adjacent in the heatmap and a line was drawn below to connect them. Insertions detected in leaves are indicated to the right, with the leaf of first detection listed in parentheses; insertions without a leaf listed were not detected in any leaf and likely arose after the leaves diverged from the tassel. Bottom, lineage reconstruction for this plant (expanded from **Fig. 2C**, top and **Fig. 3A**). Dots represent insertions in a given leaf and branch. Numbers inside the dots indicate the number of insertions that occurred at that time and branch point (if greater than 1). If the dots and internal numbers are added up for a given branch, it will match the number of insertions visible in the heatmap. **Supplementary Notes 1** and **2** describe how lineage phylogenies and precursor cell numbers were inferred.

**Fig. S6.**
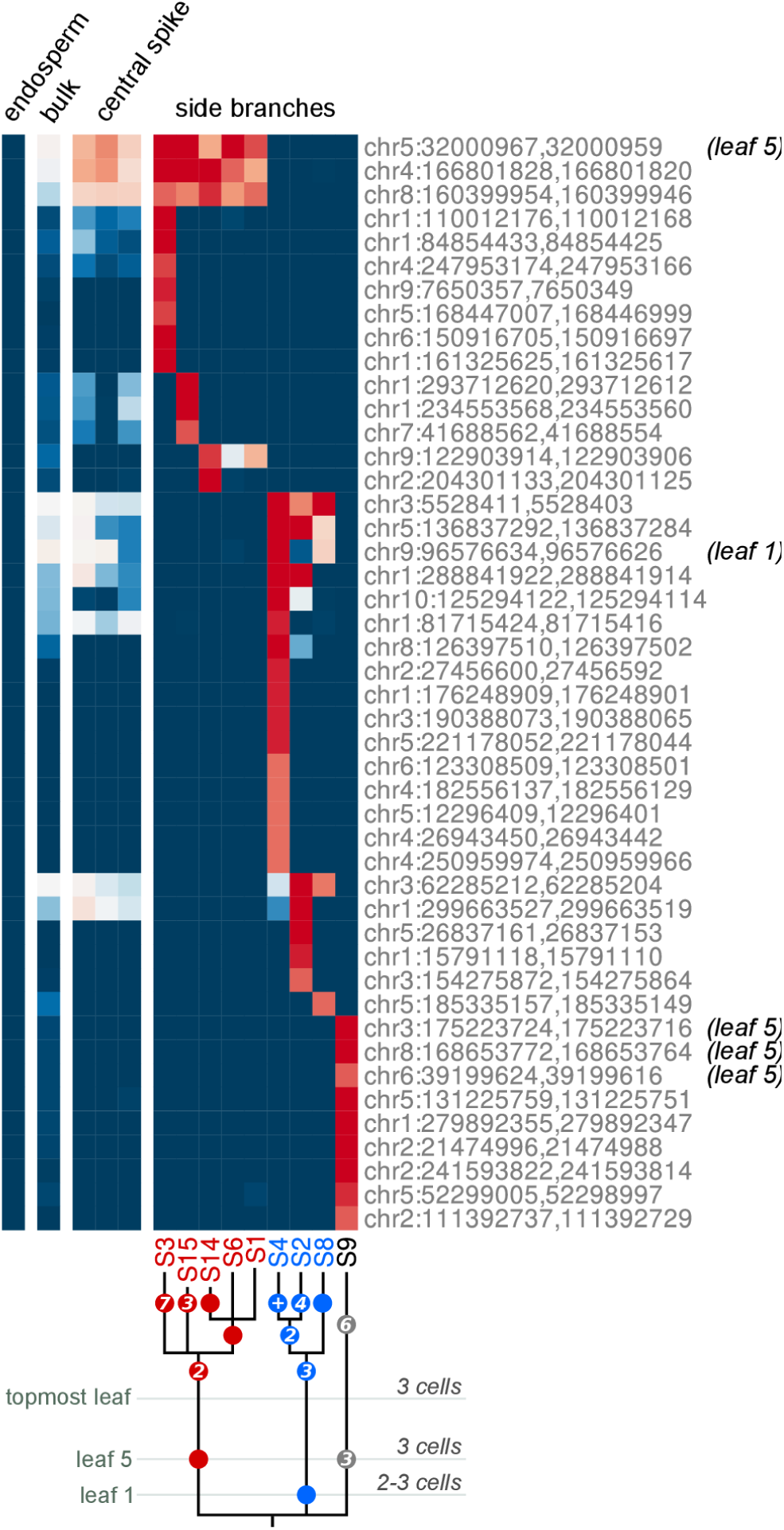
Heatmap of abundant *de novo* insertions and lineage reconstruction for plant NF51-1. This plot is the same as for **Fig. S5**. For this plant, only leaves 1, 5, 13, 15, and 16 were collected (out of 16 total leaves).

**Fig. S7.**
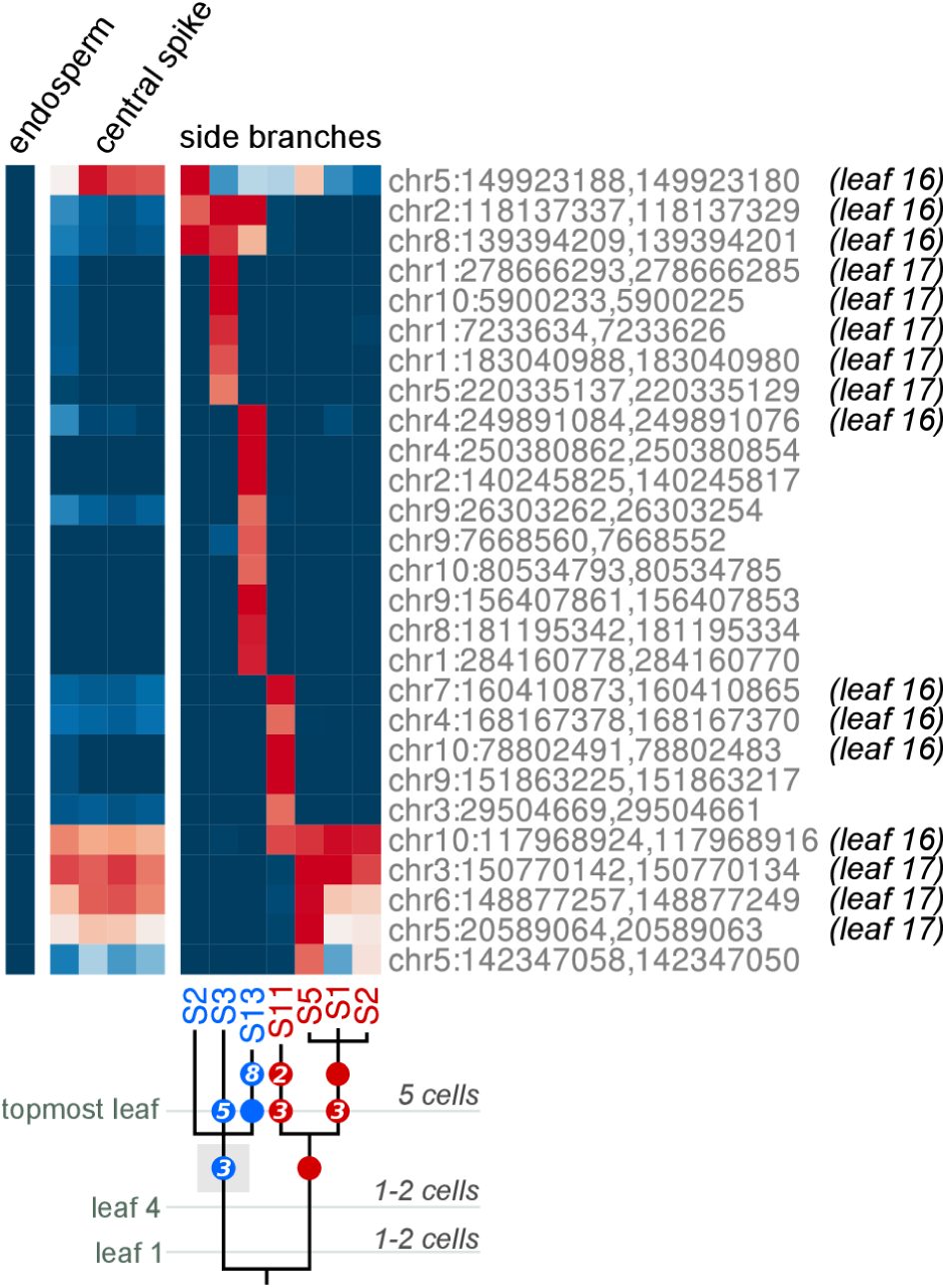
Heatmap of abundant *de novo* insertions and lineage reconstruction for plant NF51-3. This plot is the same as for **Fig. S5**. For this plant, only leaves 1, 4, 16, and 17 were collected (out of 17 total leaves).The light gray box in the lineage diagram (behind a blue dot with the number ‘3’) indicates that there was discordance between the data and lineage reconstruction; the discordance was caused by the very top insertion in the heatmap, as it was detected in every tassel branch, but at low abundance in many of them. The low abundance of this single insertion in the ‘red’ side branches is difficult to reconcile with a clean cell phylogeny, leading to uncertainty in the reconstruction. This and plant NF55-1 (**Fig. S10**) are the only two cases where any such discordance was observed (all other plants could be cleanly explained by the phylogenetic reconstruction).

**Fig. S8.**
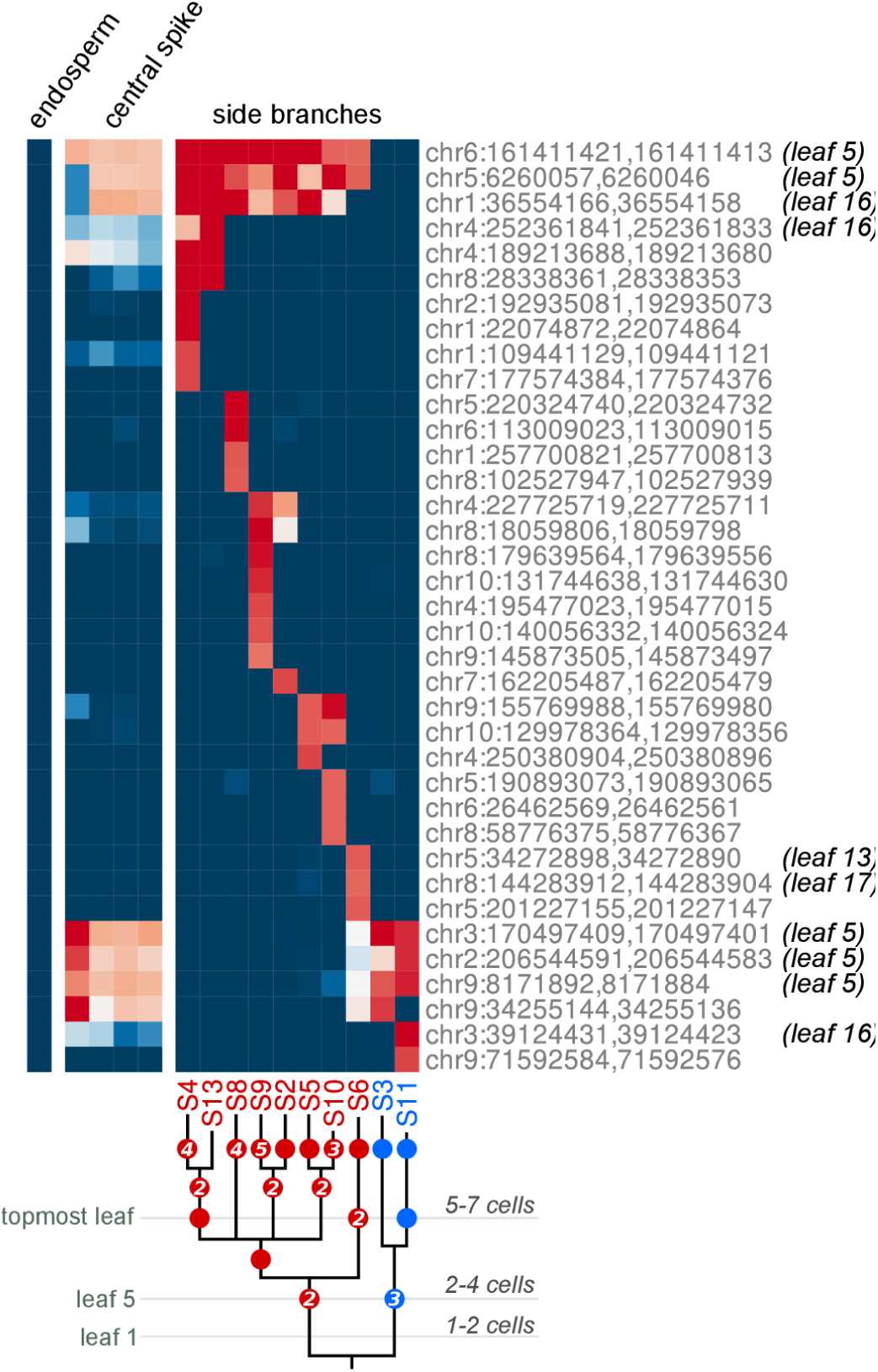
Heatmap of abundant *de novo* insertions and lineage reconstruction for plant NF51-4. This plot is the same as for **Fig. S5**. For this plant, only leaves 1, 5, 16, and 17 were collected (out of 17 total leaves).

**Fig. S9.**
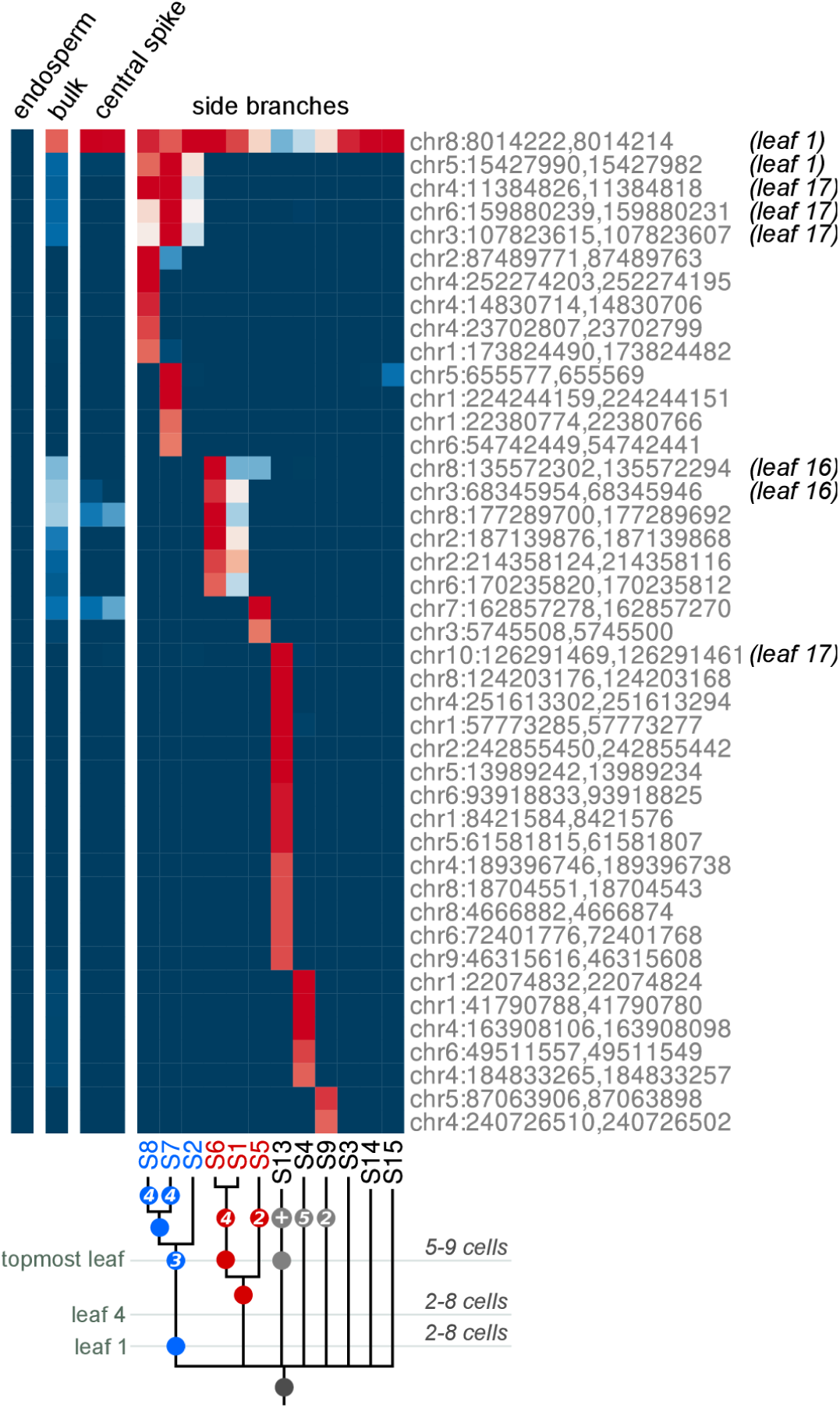
Heatmap of abundant *de novo* insertions and lineage reconstruction for plant NF53-3. This plot is the same as for **Fig. S5**. For this plant, only leaves 1, 4, 16, and 17 were collected (out of 17 total leaves). The ‘+’ in the gray dot of the lineage diagram indicates that ≥10 insertions occurred in this branch.

**Fig. S10.**
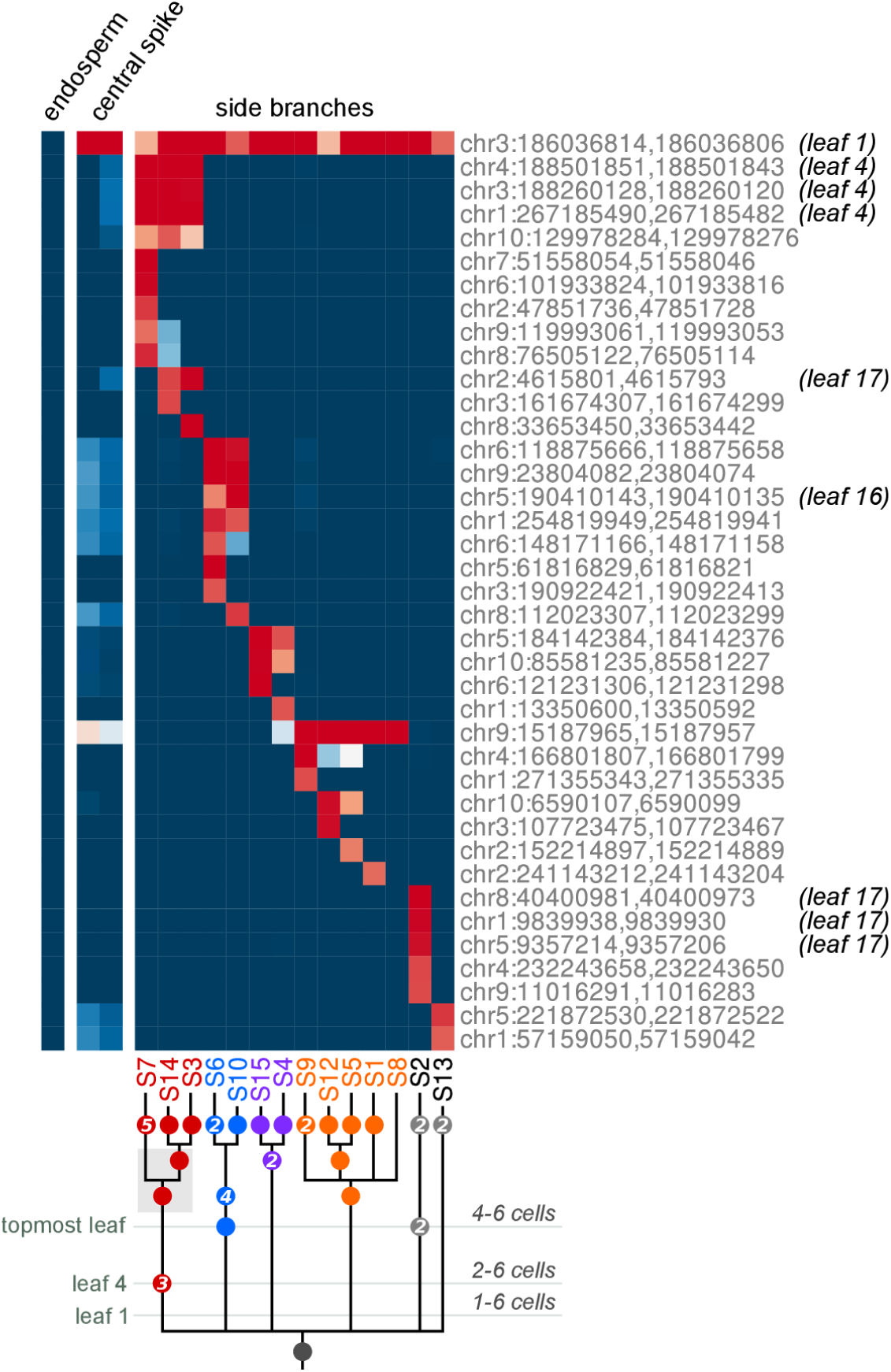
Heatmap of abundant *de novo* insertions and lineage reconstruction for plant NF55-1. This plot is the same as for **Fig. S5**. For this plant, only leaves 1, 4, 16, and 17 were collected (out of 17 total leaves). The light gray box in the lineage diagram (within the ‘red’ cluster to the left) indicates that there was discordance at that part in the lineage reconstruction. The reason is that the side branch abundance vs presence in leaves suggests a conflicting timing for these two insertions: from the side branches, one of these mutations (on chr10) was found in S7, S14, and S3, while the other (on chr2) was only in S14 and S3; this suggests that the chromosome 10 insertion occurred first, there was a cell division, and then the chr2 insertion occurred (this is how the tree is drawn). However, the putative later insertion (on chr2) was detected in the top leaf, while the earlier insertion (on chr10) was not. This is the only case where the leaf data suggests a conflicting order with the side branches. Overall, the uncertainty does not affect many conclusions. It is possible the chr10 mutation was just missed in leaves, in which case the discordance would go away, the branching would be estimated before the topmost leaf (rather than after, as currently drawn), and the number of cells at the topmost leaf would increase by 1 (5-7 cells instead of 4-6).

**Fig. S11.**
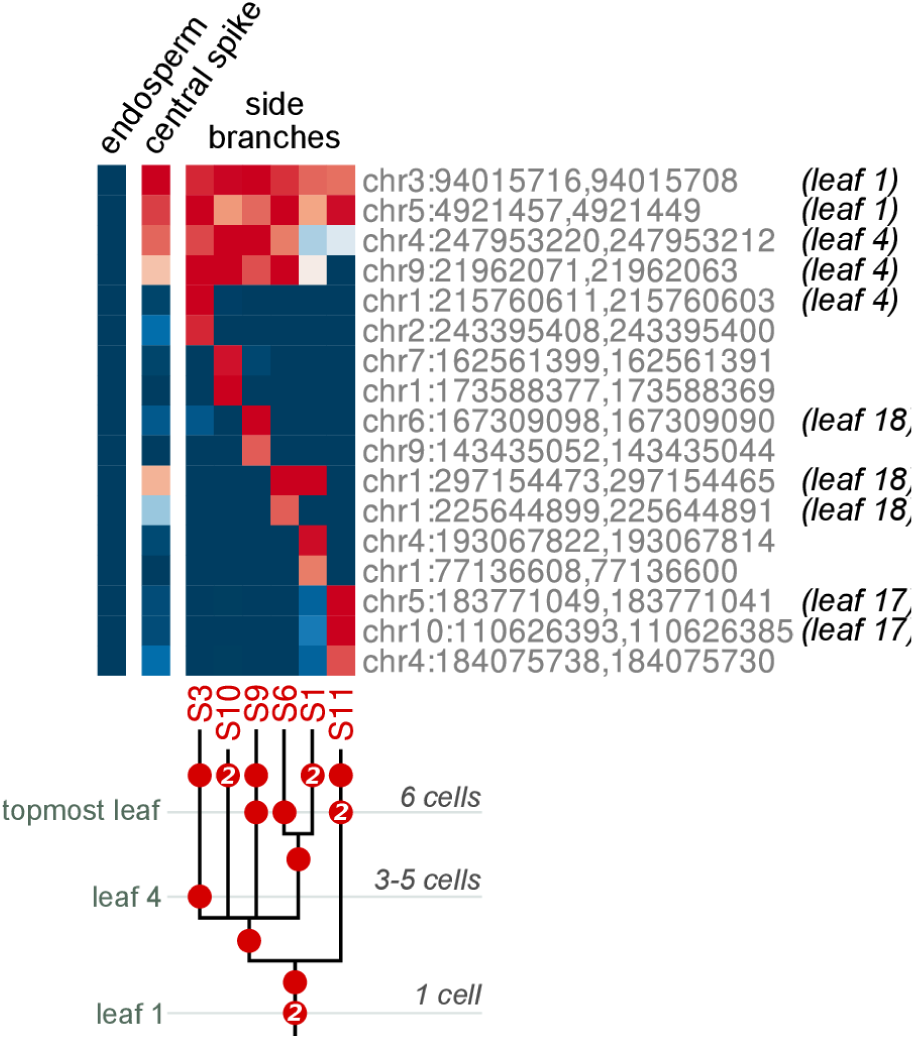
Heatmap of abundant *de novo* insertions and lineage reconstruction for plant NF55-2. This plot is the same as for **Fig. S5**. For this plant, only leaves 1, 4, 17, and 18 were collected (out of 18 total leaves). This is the only plant where there is unambiguously a single cell bottleneck at the time the first leaf diverged from the tassel. By the 4^th^ leaf, the pollen progenitor population has already expanded to at least 3 cells.

**Fig. S12.**
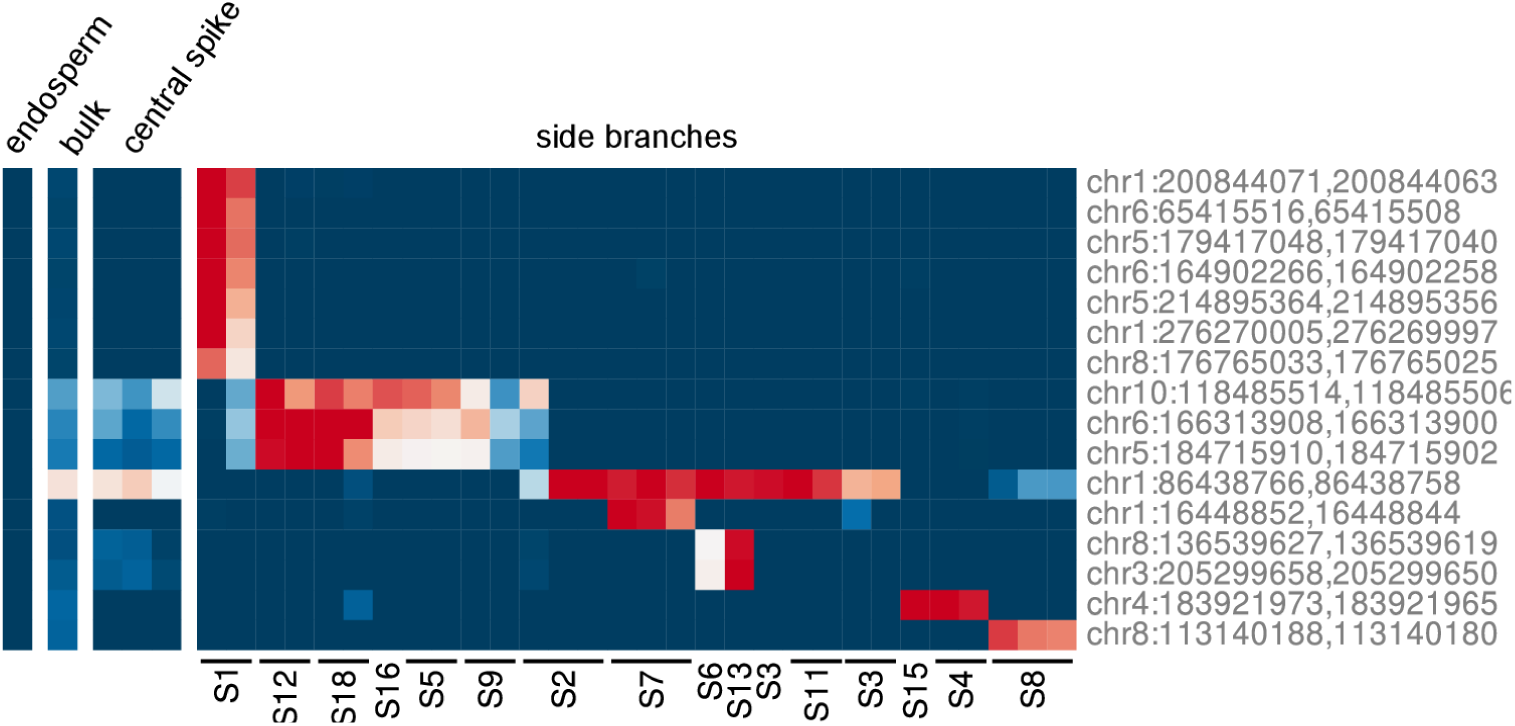
Heatmap of abundant *de novo* insertions for plant ND72-21. This plot was prepared the same as for **Fig. 2C** and **S5** (top). For this plant, no leaf tissue was collected and so the timing of insertions based on leaf number is not labeled. Several branch were collected repeatedly across different days; these repeat collections are labeled with a line as in Fig. S5. For side branches, the lineage composition was sometimes variable between days (e.g. side branch 2 above); in such cases, the branches were always located near the boundary of the respective lineages on the tassel.

**Fig. S13.**
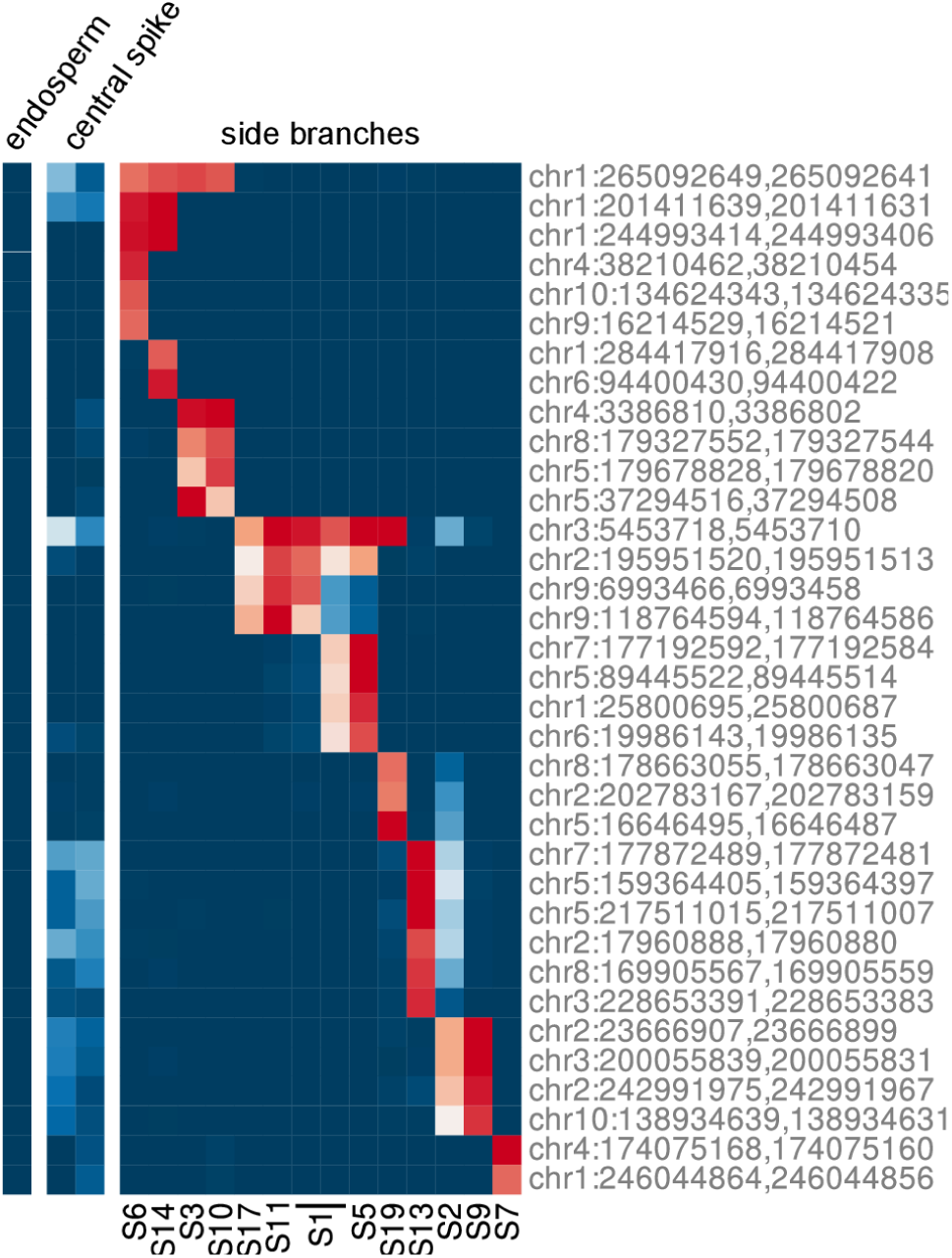
Heatmap of abundant *de novo* insertions for plant NE56-3. This plot was prepared the same as for **Fig. S12**. For this plant, no leaf tissue was collected and so the timing of insertions based on leaf number is not labeled.

**Fig. S14.**
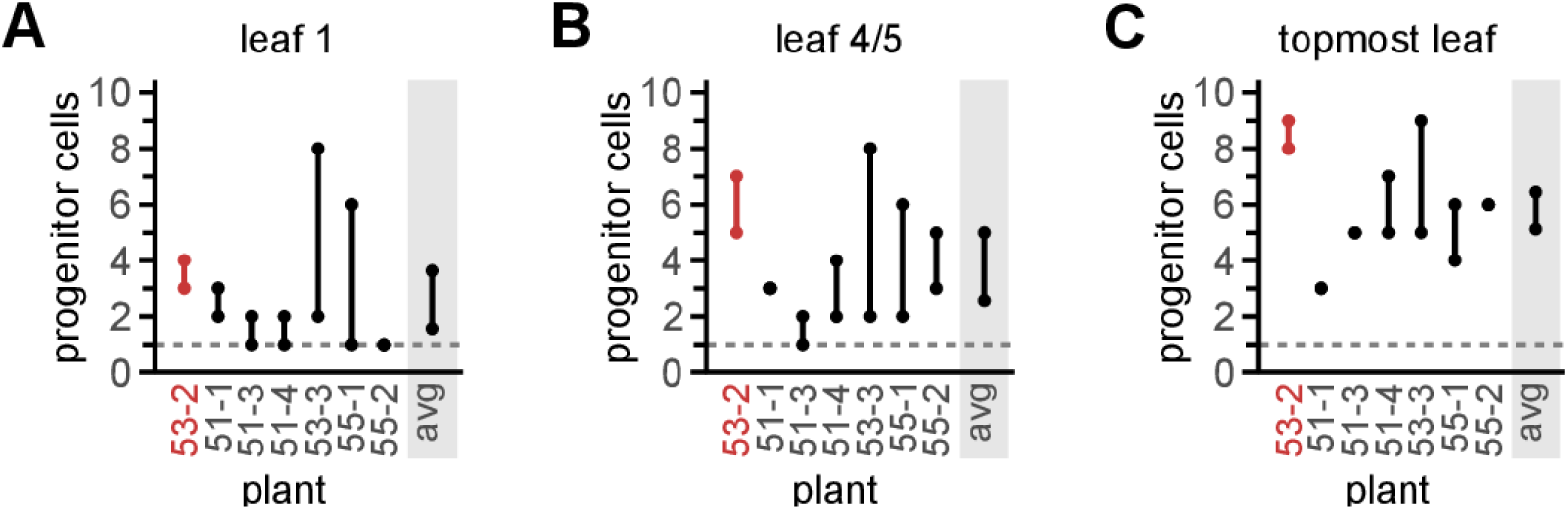
Estimated progenitor cell number leading to pollen at different stages. Estimated number of progenitor cells leading to pollen at the time of divergence from leaf 1 (**A**), leaf 4 or 5 (**B**; whether leaf 4 or 5 was collected varied by plant), or the topmost leaf (**C**). For each plant, the line shows the range of progenitor cell numbers that are consistent with the data, based on lineage reconstructions (**Fig. S5-S11**). For example, for plant 53-2 (highlighted in red as this is the plant featured in the main text), 3 or 4 progenitor cells at the time of first leaf divergence would be equally consistent with the TE insertion data. Even with ambiguities from lineage reconstruction, the data show that not all plants have an identical number of progenitors at a given stage. For instance, plant 53-2 must have at least 8 progenitors at the time the top leaf diverged (panel **C**), while plant 51-1 has only 3. This is consistent with prior conclusions that maize development is probabilistic and there is not a fixed number of cells dedicated to the tassel at given stages (*27*). Overall, the estimated number of progenitor cells increased over time during embryo development.

**Table S1.** Clonal diversity in animal germlines. With the exception of *C. elegans*, all other animal species considered have polyclonal germlines – with subsets of PGCs more related to subsets of somatic cells than each other. This would maintain lineage diversity and reduce the mutation recurrence rate, similar to the process described in maize. PGC, progenitor germ cell. *The population size is defined as the minimum PGC population after the last PGC has fully diverged from any somatic lineages. For instance, in the lineage cartoon shown in **Fig. 1A** (top), there would be a single cell at commitment, with the last common ancestor of all PGCs occurring two divisions after the zygote. Alternatively, for the cartoon in **Fig. 1A** (bottom), there would be 4 cells at germline commitment with a last common ancestor of the zygote. **Lineage tracing in the early drosophila embryo (the first 7 syncytial divisions) has not been possible by either live-microscopy or sectoring (*48*). As a result, the last common ancestor of all PGCs in *Drosophila* is unknown.

|  | # cell divisions at<br>germline<br>commitment | Population size at<br>commitment*<br>(# PGCs) | Last common<br>ancestor of all<br>PGCs | Reference<br>number(s) |
| --- | --- | --- | --- | --- |
| **Human** | 16 | unknown, polyclonal | variable, zygote | 1-3, 46 |
| **Mouse** | 11-13 | 45 | variable, zygote | 34-36, 46 |
| **Zebrafish** | 10 | 4 | zygote | 33 |
| ***C elegans*** | 4 | 1 | P4 | 47 |
| ***Drosophila*** | 10 | 8-10 | unknown** | 36 |

**Supplementary Note 1**

This Supplementary Note walks through the process used to infer pollen lineage phylogenies from Mu TE insertion data (**Fig. 2C**, **Fig. 3**, and **Fig. S5-S11**). These phylogenies were further used to plot spatial distributions in **Fig. 2D-E** and estimate the number of pollen progenitor cells during different stages of development (see **Supplementary Note 2**).

Cell phylogenies were inferred from the presence or absence of TE insertions in pollen from different tassel branches. For example, for the plant in **Fig. 2** there were five *de novo* TE insertions abundant in side branches 3, 5, 8, 11, 12, and 13 but largely absent from all others:

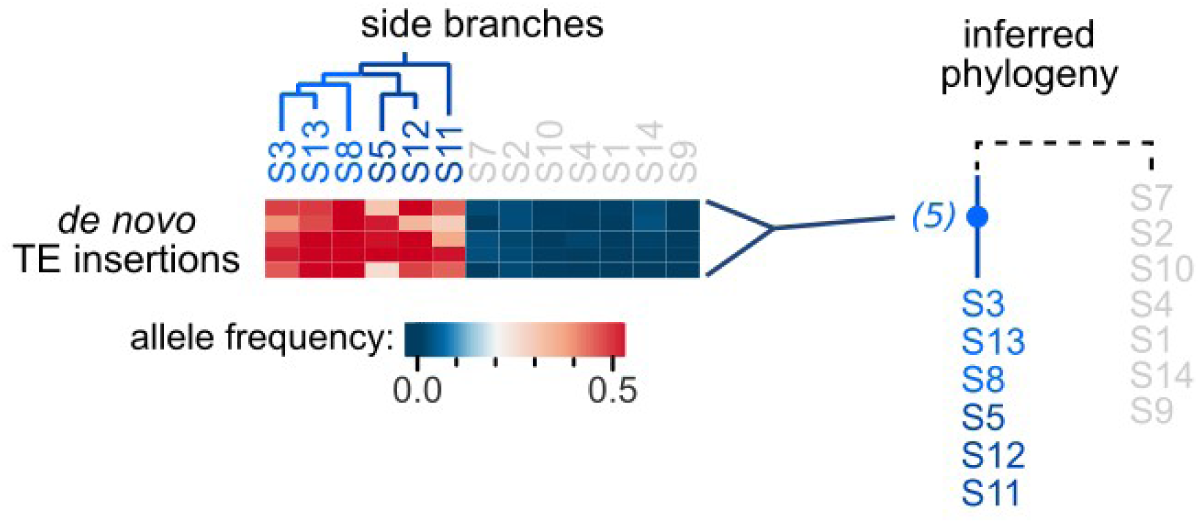

This suggests that these side branches were primarily derived from the same progenitor cell that contained the 5 insertions, while the other side branches were derived from a distinct cell or cells. In **Fig. 3**, matched leaf data provides further resolution, as two of these five insertions were present starting from the first seedling leaf, while the other three were detectable by leaf 2:

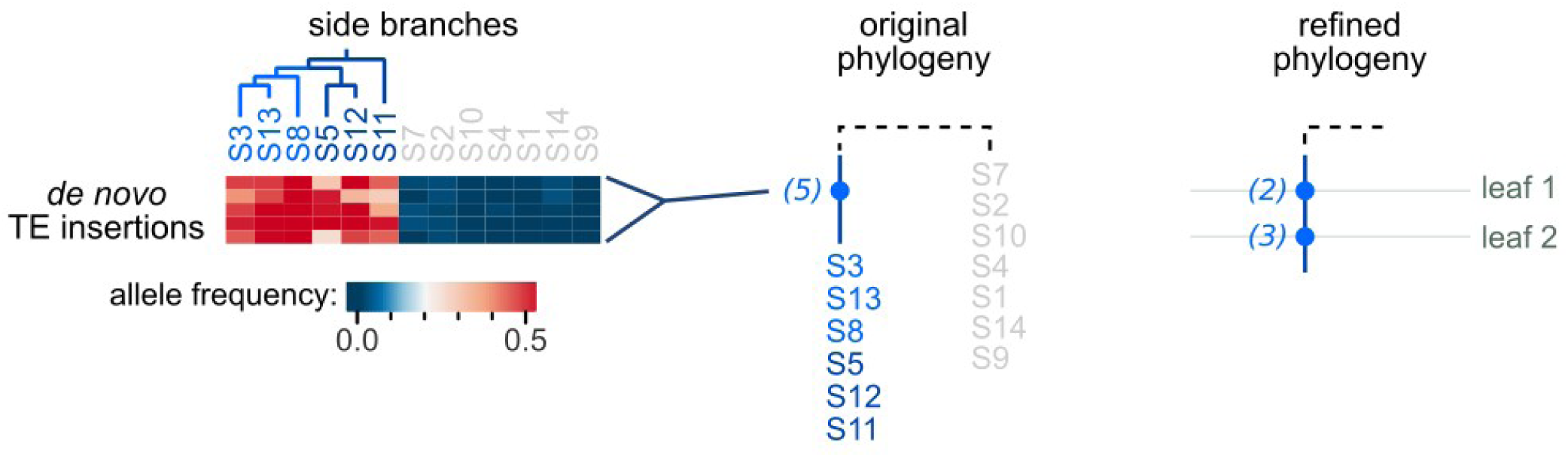

In this case, the leaf data provides more refined timing of the events. First, it indicates two of the insertions likely occurred earlier than the other three. Second, by comparison to sectoring studies (*27*), insertions shared with leaf 1 likely occurred very early in the embryo before even layer 1 (L1) and layer 2 (L2) diverged. Similarly, the three insertions shared with leaf 2 likely arose before the meristem had formed, as X-ray induced sectors spanning from leaf 2 to the tassel were only observed when treating embryos at earlier stages (*27*).

The next two TE insertions were present in only five of the six ‘blue’ side branches, suggesting there had been a split into at least 2 cells at the time these insertion occurred: one cell with these two insertions gave rise to pollen from five of the branches, while the cell(s) without the insertions contributed to pollen from ‘S11’. In this case, both insertions were first detected in leaf 5, helping again to refine the timing of events:

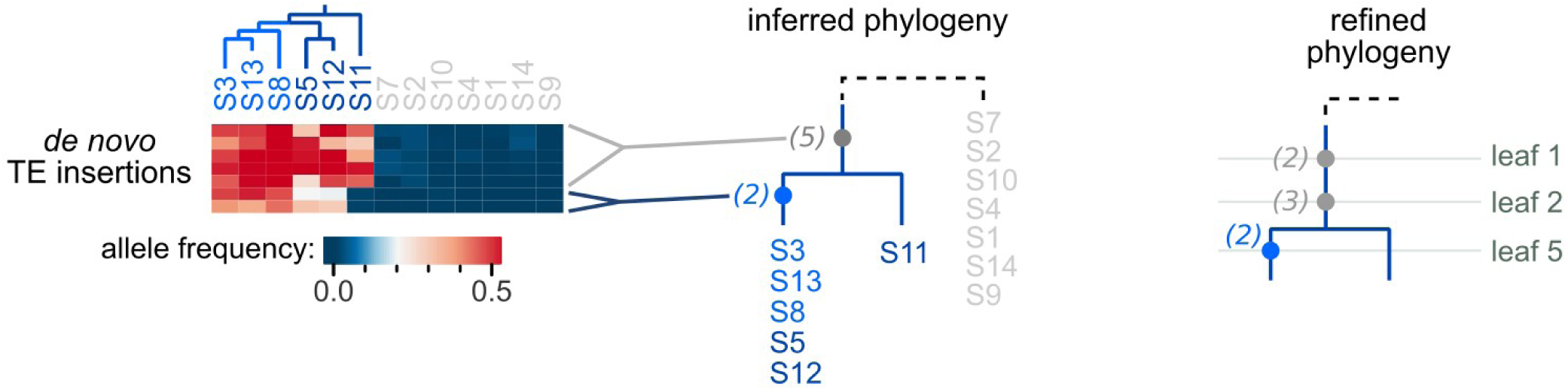

In the figure above and following figures, new parts of the phylogenetic reconstruction are shown in blue, while previously discussed segments are in gray.

Next, two TE insertions were observed in S3, S13, and S8 but not the other side branches, suggesting these three branches were descended from a single common ancestor more recent than S5 or S12:

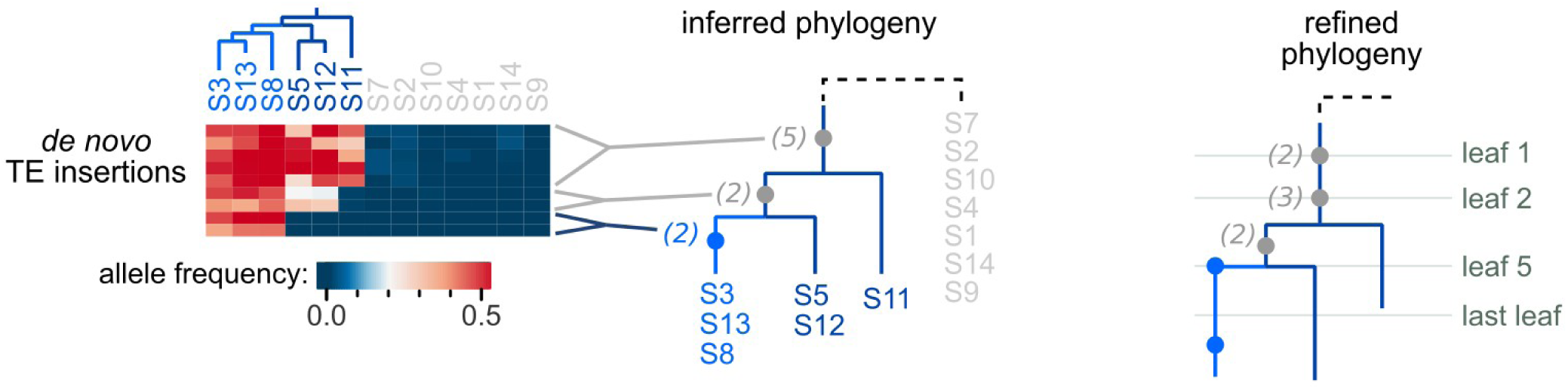

One of these insertions was present in leaf 5, the same leaf in which the previous two insertions were first observed. This is an example where the leaf timing provides less resolution than the initial phylogeny: we can infer that the insertions present in S5 and S12 likely occurred first, even though one of the newer insertions was identified in the same leaf (leaf 5). The second insertion in S3/S13/S8 was not detected in any leaves and likely occurred much later. There were several cases like this, where sometimes the matched leaves provided greater resolution about the order of events and other times the phylogenetic inference had greater resolution.

Here is a circumstance where ambiguity exists in the phylogeny. There were no insertions that connect side branch 5 and 12 with each other or with the ‘light blue’ group (S3, S13, S8)… this means there is uncertainty about the relationship between S5, S12 and the other side branches – three equally parsimonious phylogenies can be drawn from the data:

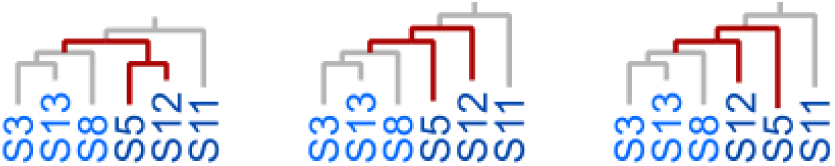

In the main text, we show the phylogeny on the left as the most probable (with S5 and S12 sharing a lineage), because these two side branches are located directly above each other along the tassel circumference and so this is more likely based on spatial location (**Fig. 2D**). However, the mutation data is fully consistent with the other two phylogenies.

Finally, five additional insertions indicate that S3 and S13 share a common ancestor distinct from S8. All five of these insertions were absent from all leaves, suggesting they arose in the tassel primordia after leaf divergence:

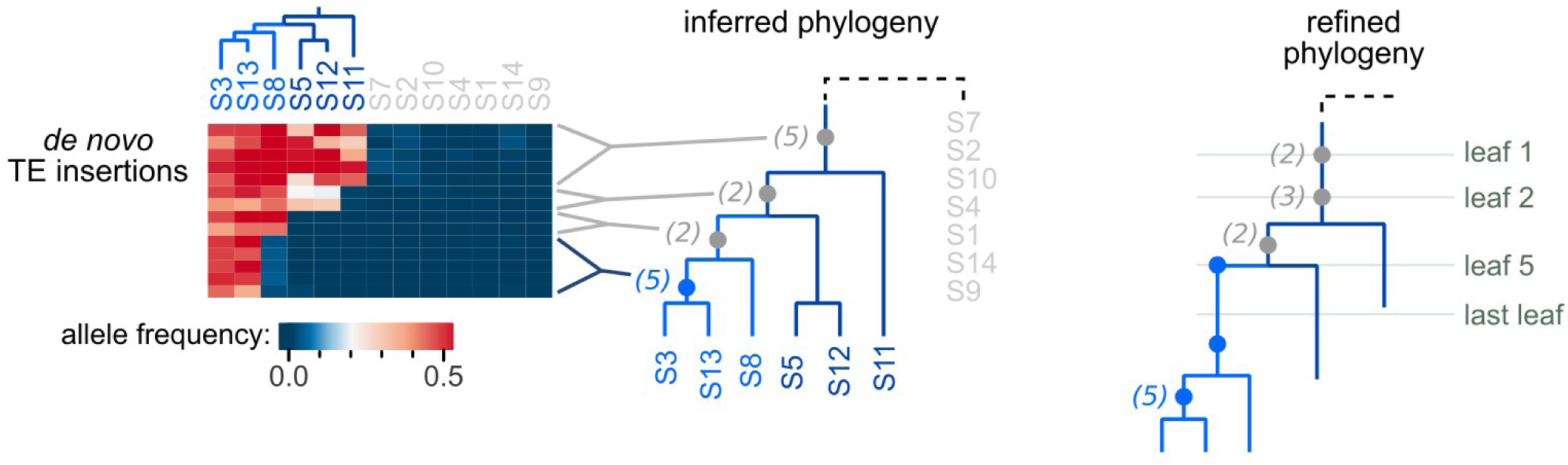

This completes the phylogenetic reconstruction for the ‘blue’ group in **Fig. 2C** and **3A**. The ‘inferred phylogeny’ above is the same as what is shown about the heatmap in **Fig. 2C**. The ‘refined phylogeny’ is the same as what is shown in both **Fig. 3A** (which focuses on the parts of the phylogeny before the last leaf) and **Fig. S5** (which also includes the phylogeny beyond the last leaf). Similar logic applies to the remainder of the phylogenetic reconstructions.

**Supplementary Note 2**

The number of pollen progenitors at the time a given leaf diverged can also be estimated from the phylogenetic reconstructions and TE insertion data. For instance, for the plant shown in **Fig. 3**, the ‘blue’ lineage must have descended from one (and exactly one) cell at the time the pollen progenitors diverged from leaf 1. This can be inferred from five TE insertions that were present in every side branch of the ‘blue’ lineage:

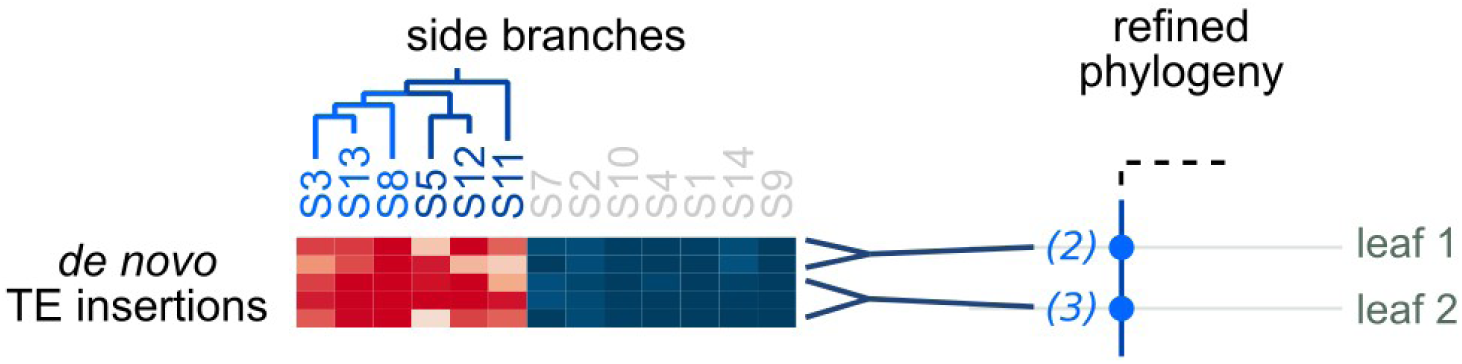

Because two of these insertions were detected in leaf 1, this lineage must have already formed by the time leaf 1 diverged in the embryo. This defines the minimum number of cells contributing to the blue lineage.

To define the maximum number of possible progenitor cells, we must look beyond leaf 1. In this case, there were three insertions shared by leaf 2 that were found in all six ‘blue’ side branches. The presence of these insertions in leaf 2 indicates that, sometime after leaf 1 divergence, there still must be only one cell contributing to this lineage. Thus, the maximum number of progenitor cells contributed by the blue lineage is one.

For clarity, consider some alternative scenarios… if all five of these insertions were found in leaf 1, then it would be possible for either one or two cells to contribute to the phylogeny:

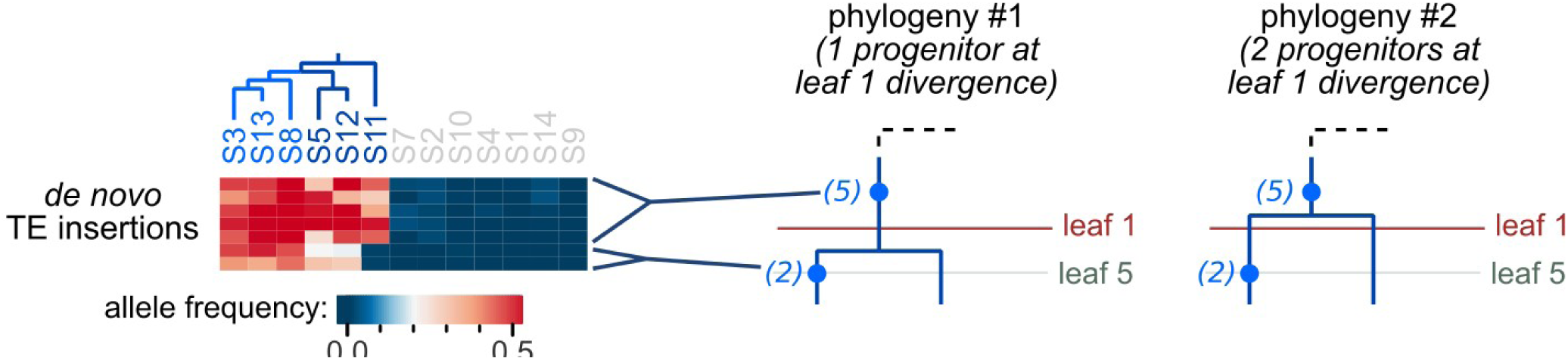

Thus, the insertions shared within a leaf help define the minimum number of progenitors consistent with the data, while it is critical to consider insertions in subsequent leaves to determine the maximum number.

In total for this plant, there can be 3 or 4 progenitor cells (see **Fig. 3A**): exactly 1 progenitor from the blue lineage (discussed above), exactly one from the purple lineage (for similar reasons), and either 1 or 2 from the red and orange lineages – there were no TE insertions shared with the first leaf that marked the red or orange lineages, so it is possible that these lineages descended from a single cell progenitor at the time of leaf 1 divergence.

